# Novel insights into Emx2 and Dmrta2 cooperation during cortex development and evidence for Dmrta2 function in choroid plexus

**DOI:** 10.1101/2024.09.19.613943

**Authors:** Jithu Anirudhan, Xueyi Shen, Tünde Szemes, Marc Dieu, Abdulkader Azouz, Louise Conrard, Gilles Doumont, Maren Sitte, Younes Achouri, Sadia Kricha, Gabriela Salinas-Riester, Patricia Renard, Eric J. Bellefroid

## Abstract

Early dorsal telencephalon development is coordinated by an interplay of transcription factors that exhibit a graded expression pattern in neural progenitors. How they function together to orchestrate cortical development remains largely unknown. The *Emx2* and *Dmrta2* genes encode TFs that are expressed in a similar caudomedial^high^/ rostrolateral^low^gradient in the ventricular zone of the developing dorsal telencephalon with, in the medial pallium, *Dmrta2* but not *Emx2* expressed in the developing choroid plexus. Their constitutive loss has been shown to impart similar cortical abnormalities, and their combined deletion exacerbates the phenotypes, suggesting possible cooperation during cortex development. In this study, we utilized molecular and genetic approaches to dissect how Emx2 functions with Dmrta2 during cortical development. Our results show that while they regulate a similar set of genes, their common direct targets are limited but include key regulators of cortical development. Identification of the interaction partners of Emx2 suggests that it coordinates with the LIM-domain binding protein Ldb1 to execute the activation and repression of some of its downstream targets. Finally, while *Emx2* is known to suppress choroid plexus development, we also provide evidence that *Dmrta2* is in contrast required for choroid plexus since in its absence in medial telencephalic progenitors, mice develop hydrocephalous postnatally, a phenotype that appears to be due to a compromised cytoarchitecture. Together, these data indicate that Emx2 and Dmrta2 have similar but also distinct functions in telencephalon development and provide the first insights into Emx2 mechanism of action.

**SIGNIFICANCE STATEMENT:** *Emx2* and *Dmrta2* encode transcription factors that generate similar phenotypes upon their loss in the developing cortex suggesting possible cooperation. Here we explored how Emx2 functions with Dmrta2 during cortical development. Results obtained indicate that Emx2 directly regulates with Dmrta2 only a few genes, some coding for key cortical determinants and that Emx2 utilizes the Ldb1 cofactor for the regulation of some of its targets. Results also suggest that, unlike Emx2 which suppresses choroid plexus development, Dmrta2 is required for choroid plexus as its loss in medial telencephalic progenitors leads to hydrocephalus. Together, our results reveal that Emx2 and Dmrta2 have similar but also distinct functions during telencephalon development and provide novel insights into the mechanism of action of Emx2.

## INTRODUCTION

Early cerebral cortex development is orchestrated by morphogens produced by distinct signaling centers and transcription factors (TFs) gradients in dorsal telencephalic progenitors. Their action has far-reaching implications for adult animals’ cortical architecture and behaviour (Leingärtner et al., 2007; O’Leary et al., 2007; Hébert & Fishell, 2008; Hoch et al., 2009). The key telencephalic organizers that set up the transcription factor gradients across the cortical neuroepithelium include the *Fgf8* expressing Anterior Neural ridge (ANR), the Wnt and BMP expressing cortical hem, the Wnt antagonist rich anti-hem and the Shh expressing ventral telencephalon (Agirman et al., 2017; Grove & Monuki, 2020). Several transcription factors (TFs) that exhibit a graded expression have been shown to be important in early cortical development. Among them, the homeobox *Emx2* and the zinc finger *Dmrta2* TFs, whose expression starts immediately after the specification of the cortical primordium, are expressed in a similar high caudomedial to a low rostrolateral gradient in dorsal telencephalic progenitors. However, in the medial pallium, *Dmrta2* but not *Emx2* is expressed in the developing choroid plexus (ChP) suggesting dissimilar function in this tissue (Tole et al., 2000; Konno et al., 2012; Saulnier et al., 2013; Ypsilanti et al., 2021).

*Dmrta2* and *Emx2* single knockouts impart similar phenotypes, both leading to a reduction of caudo-medial cortical structures. Indeed, in *Dmrta2* knockout embryos, the cortical hem is strongly reduced, leading to the loss of the hippocampus and the dentate gyrus. In *Emx2* knockout, the hippocampus is shrunken, with no structurally identifiable dentate gyrus. In *Emx2*;*Emx1* double knockout, a more severe phenotype occurs with loss of the hippocampus and dentate gyrus (Yoshida et al., 1997; Tole et al., 2000; Saulnier et al., 2013). In both *Dmrta2* and *Emx2* single knockouts, concomitant with the loss of medial structures, there is a reduction in overall cortical size and an expansion of the paleocortex. Both single mutants also exhibit lamination defects, TCA innervation defects and reduced Cajal-Retzius cell number (Mallamaci et al., 2000; López-Bendito et al., 2002; Mallamaci, 2011; Konno et al., 2012; Saulnier et al., 2013; Ratié et al., 2020). Both TFs have been also shown to contribute to the control of neurogenesis and the suppression of astrogliogenesis (Falcone et al., 2015; Muralidharan et al., 2017). In addition, both influence the arealization of the neocortex in a dose-dependent manner. Indeed, the loss of either of them leads to a reduction of caudal areas and an expansion of rostral areas (Bishop et al., 2000, Saulnier et al., 2013). By contrast, either of their over-expression creates an opposite arealization phenotype (Hamasaki et al., 2004, De Clercq et al., 2018). Interestingly, their compound deletion leads to an even more dramatic reduction in cortical size at E12.5. Our previous work has also shown that Dmrta2 and Emx2 both bind and regulate the expression of *Gsx2* in the pallial-subpallial boundary through their adjacently placed motifs in a distal enhancer (Desmaris et al., 2018). The similarity of their expression and functions, and the exacerbation of the phenotype in the event of their combined loss, points to their possible cooperation during cortical development.

Though genetic studies have uncovered the crucial role played by many TFs in the context of cortical patterning and development, molecular mechanisms that distinct TFs utilise for their combinatorial outputs during cortical development are only beginning to be uncovered. In this study, to try to understand how Emx2 functions in cooperation with Dmrta2 to control cortex development, we performed bulk-RNA analysis on *Emx2* KO embryos and compared their global gene expression to that of *Dmrta2* KO embryos, and searched for Emx2 direct targets and interacting partners. Besides, speculating that Dmrta2 may have a unique function in relation to Emx2 in choroid plexus development, we also examined the role of *Dmrta2* in this tissue. Our results provide novel insight into the Emx2 mechanism of action in cortex development and reveal a novel function for *Dmrta2* in choroid plexus tissue.

## MATERIAL AND METHODS

### Mice

All animal procedures were approved by the CEBEA (Comité d’éthique et du bien-être animal) of the IBMM-ULB and conformed to the European guidelines on the ethical care and use of animals. The mid-day of the vaginal plug appearance was defined as embryonic day 0.5 (E0.5) for timed pregnancies. Mice were provided *ad libitum* with standard mouse lab pellet food and water and housed at room temperature (RT) with a 12h light/dark cycle. All mice were maintained either in a C57BL/6 or CD1/C57BL6 mixed background, and litter mate controls of either sex were used in experiments. For *Lmx1a*-Cre animals, sex genotyping was performed, and only male embryos were used for experiments since the Cre line was X-linked, and Cre-positive adult females were used to maintain the line as recombination was observed in male testis.

Mouse lines previously described include *Dmrta2^+/-^* (Saulnier et al., 2013), *Dmrta2^fl/fl^*(De Clercq et al., 2018), *Emx2^+/-^* (Pellegrini et al., 1996), *Lmx1a-Cre* (Chizhikov et al., 2010), *Rosa-Stop-EYFP* (Srinivas et al., 2001), *Msx1:Cre^ERT2^*(Lallemand et al., 2013). *Emx2^+/-^* mice was a kind gift from Dr. Antonello Mallamaci, SISSA, Italy. *Lmx1a-Cre* mice was a kind gift from Dr. Kathleen J Millen, University of Washington, USA. *Rosa-Stop-EYFP* and *Msx1:Cre^ERT2^* lines were a kind gift from Dr. Michel Cayouette, IRCM, Canada.

The *Emx2* knockin (Emx2 KI) mouse line was generated using the cloning-free CRISPR-Cas9 system (Aida et al., 2015). Guide RNAs were designed using CRISPR direct (Naito et al., 2015). A 20 nucleotides crRNA targeting the c -terminal end of *Emx2* ORF, (5’-TTTGGACTTTTAATCGTCTG-3’, Integrated DNA Technologies) corresponding to the target sequence with a tail, a 67 nucleotides tracRNA (ref 1072534, Integrated DNA Technologies) containing the hairpin recognised by Cas9 with an extremity complementary to the crRNA tail) and a long single-stranded oligonucleotide containing homology arms to the targeted region and a 3xFlag-V5 sequence (Megamer® ssDNA Fragment, Integrated DNA Technologies: 5’CCACATTAACCGGTGGAGAATTGCTACCAAGCAGGCGAGTCCGGAGG AAATAGATGTGACCTCAGACGATGACTACAAAGACCATGACGGTGATTATAAAG ATCATGACATCGACTACAAGGATACGATGACAAGGGCTCGGGCTCGACCTCGGG TAAGCCTATCCCTAACCCTCTCCTCGGTCTCGATTCTACTAAAAGTCCAAACCCA TCCTACAAAAATGGACAGCACAGAGCAAGAAAGACAGGGAGAGGAGGGAGAAA A -3’) were synthesised. After annealing the crRNA/tracrRNA (2.4 pmol/μl), recombinant Cas9 nuclease V3 (200 ng/μl; Integrated DNA Technologies) and the single-strand DNA donor (5 ng/μl each) were mixed in IDTE buffer (11-01-02-02, Integrated DNA Technologies) and injected into the pronucleus of B6D2F1 mouse zygotes. Injected zygotes were transferred into oviducts of CD1 pseudo-pregnant female mice after incubating them in KSOM medium at 37°C for at least 2 hours. Positive founders were identified by genotyping using primers Fwd– 5’ GAG GAA GAA GGC TCA GAT TCT 3’; Rev – 5’ CCTCGGTCTCGATTCTACATAA 3’)followed by sequencing. Two founders were selected to establish the colonies after confirming the expression of tagged protein through Western Blotting using lysate derived from knockin mice cortices.

For tamoxifen administration, tamoxifen dissolved in corn oil was administered to timely pregnant dams intraperitoneally on the E9.5 day of pregnancy for 75mg/kg of body weight of mice. Dams were sacrificed on the E12.5 day of pregnancy.

### Cell lines

P19 cells (ATCC*-*CRL*-*1825) or HEK293T cells (ATCC-CRL-3216) were maintained in DMEM medium supplemented with 10%FBS (Gibco, Cat# 26140079), and 100 U/ml penicillin-streptomycin (15140-122, Gibco) at 37°C at 5 % CO2 concentration. Transfections were carried out when cells reached 70% confluency using CalPhos Mammalian Transfection Kit (Takara Bio, Cat# 631312).

### Plasmids

The HA-Flag-Ldb1 construct was a kind gift from Dr. Nicoletta Kessaris. HA-RBBP4 plasmid was a kind gift from Dr. Terry Magnuson. pCS2-Flag-Emx2, pCS2-Emx2-3xFlag-V5 and pCS2-Flag-Emx2-3xFlag-V5, pCS2-Flag-Dmrta2, pCS2-Flag-Dmrta2ýDM were constructed in the lab and maps are available upon request.

### Antibodies

The following primary antibodies were used: anti-V5 antibody (Abcam, Cat# ab15828), anti-GFP (Aves labs, Cat# GFP1010), anti-Flag (Sigma, Cat# F3165), anti-HA (Abcam, Cat# ab9110), anti-Ldb1 (Santa Cruz, Cat# sc-365074), anti-ZO-1 (Cat# ZO1-1A12, Thermo Fisher Scientific). The following secondary antibodies were used Anti-rabbit IgG HRP conjugated (Cellsignaling, Cat# 7074), Anti-mouse IgG HRP conjugated (Jackson ImmunoResearch, Cat# 115-035-003), Anti-mouse Alexa 594 (Invitrogen, Cat# A11032), Anti-rabbit 594 (Invitrogen, Cat# A21207), Anti-mouse Alexa 488 (Invitrogen, Cat# A21202), Anti-chicken Alexa Fluor 488 (BioConnect, Jackson, Cat# 313-545-003). Normal Rabbit IgG (Cell Signaling, Cat# 2729), Anti-Digoxigenin AP, Fab Fragments (Roche, Cat# 11093274910). Normal Mouse IgG (Sigma, Cat# 12-371).

### *In situ* hybridization

Brains were harvested in ice-cold RNAse-free PBS and fixed overnight in 4% PFA in RNAse-free PBS. Brains were subsequently washed twice the next day in RNAse-free PBS and kept in 30% sucrose in RNAse-free PBS. Once the brains were equilibrated, they were embedded in a 15% sucrose 7.5% gelatin in RNAse-free PBS and stored at -80°C. The brains were coronally sectioned on a cryostat at 20µm thickness and collected on Superfrost (Epredia, Cat# J1800AMNZ) slides. The slides were allowed to dry for at least 20 minutes and stored at -20°C until further use. Slides were first thawed to room temperature and post-fixed in 4%PFA for 20min, washed twice in RNase free 1X PBS-0,1% Tween20 and incubated in 1X PBST with Protease K dissolved in DEPC water(10μg/ml) (Sigma Aldrich) for 3min for permeabilising the tissue. The reaction was stopped by adding in Glycine (2mg/ml) in PBS. The sections were post-fixed in 4%PFA along with glutaraldehyde (0.2%) and washed three times. Subsequently, sections were incubated in the hybridization solution (50% formamide, SSC 5X pH4,5, tRNA 25μg/ml, 1% SDS, heparin 20μg/ml) for a minimum of 3 hrs at 70°C. The sections were then incubated in appropriate RNA probes(1.5μg/mL) diluted in the same hybridization solution for overnight at 70 °C. The next day, sections were washed thrice in Solution I (50% formamide, SSC 5X pH=4,5, 1% SDS) and Solution III (50% formamide, SSC 2X pH=4,5) at -68°C and were allowed to cool down to room temperature before being washed three times in TBST 1X TBS - 0,1%Tween20. Slides were then incubated in blocking buffer (5% goat serum (Invitrogen, 10000C) diluted in TBST) for at least 2hrs at RT. Slides were then incubated with Anti-Digoxygenin fab fragments conjugated to alkaline phosphatase (Roche, Cat# 11093274910) at 1:2000 dilution overnight at 4°C. The next day, slides were washed three times in TBST and equilibrated with AP buffer (100mM NaCl, 100mM Tris pH 9, 50mM MgCl2, Tween20 0,1%). The color was revealed by incubation slides in a solution containing NBT/BCIP substrate (Roche, 4-nitroblue tetrazolium chloride, Cat# 70210625; 5-Bromo- 4- chloro-3-indolyl phosphate, Cat# 70251721). Until sufficient colour was developed, a fresh colouring solution was added to slides every 1-2 hours. Once sufficient colour developed, slides were rinsed in PBS twice and dehydrated in ethanol (70%, 90% and 100%) and HistoClear (NaAonal DiagnosAcs) and mounted using mounting EuKitt hardening mounting medium (Fluka AnalyAcal). All probes used previously described include *Emx2*, *Wnt3a, Wnt8b, Lmx1a, Lhx2, Msx1, Ttr*, *Pax6*, *Dmrta2* exon3 (Saulnier et al., 2013), *Dmrta2* exon 2 (De Clercq et al., 2018), *Emx1, Gsx2, Isl1(*Desmaris et al., 2018*), Nfib* (Hasenpusch-Theil et al.., 2012)*, Gmnc* (Arbi et al., 2016), *Nkx2.1* (Sandberg et al., 2016), *Tle3*(Chytoudis-Peroudis et al., 2018, *Tle1(*Muzio et al., 2005). Probes newly used in this study were generated from *Prdm16* (IMAGE 5704629), *Tox* (IMAGE:1428917)*, Pbx1*(IMAGE:3492658)*. Dach1* probes were generated from cDNA using primers. F 5’-GTTAGCCATCCTCCTCTCAACCATCTGC-3’, R 5’-TCATTTAAGACCCGGAGACTGTCCG-3’. Images were acquired with an Olympus SZX16 stereomicroscope and an XC50 camera, using the Imaging software CellSens.

### Immunofluorescence

Brains were harvested in ice cold PBS and fixed for 20 min in 4% PFA in PBS. Brains were subsequently washed twice in PBS and kept in 30% sucrose in 1X PBS. Once the brains were equilibrated, they were embedded in a 15% sucrose 7.5% gelatine in 1X PBS and stored at -80°C. The brains were coronally sectioned on a cryostat at 16μm thickness and collected on a superfrost (Epredia, Cat# J1800AMNZ) slides. Slides were permeabilised in 0.1% TritonX100 PBS with 3 times 15-minute wash and were blocked in 10% anti-donkey serum (Sigma, Cat# D9663) in 0.1% Triton X100 PBS for at least 2 hours. Appropriate antibodies were diluted in the blocking solution, and the slides were incubated overnight at 4°C. Subsequently, slides were washed in 0.1% TritonX100 PBS thrice for 15 minutes each and incubated with appropriate secondary antibody for 1hr at RT. The slides were washed thrice for 15 minutes and mounted using Dako Fluorescent mounting medium (Agilent, Cat# S3023). The images were taken on a Zeiss Axio Observer Z1 fluorescent microscope or a Zeiss LSM710 and were processed in FIJI.

### µPET-CT imaging

P21-30 littermate WT *(n=4)* and cKO *(n=3)* were starved overnight before intravenous injection of 4.65-5.01MBq of [^18^F]-fluorodeoxyglucose ([^18^F]-FDG) diluted in 100μL of 0.9% saline solution. Injections were done under gaseous anaesthesia (vaporized isoflurane). Following injection mice were kept anaesthesized for 10 min. Mice were then imaged on a nanoScanPETCT with Nucline v2.01 (Mediso Hungary), 2 per 2 or 3 per 3, during 15 min and starting 1h post-injection. PET was performed in 3-to-1 coincidence mode in normal count rate and reconstructed with a fully three-dimensional iterative algorithm (TeraTomo from Mediso, with 4 iterations, 6 subsets, normal regularization setting, median filtering period defined from iteration counts) to get a voxel size of 400µm (‘normal’ mode). CT imaging started automatically after the end of the associated PET for 5 min. CT images were generated for localization, attenuation, and scatter correction of PET images. CT acquisition parameters were 50 kV for a tube current of 520 μA, 300 ms per projection, 480 projections per rotation, a 4-to- 1 frame binning and a cubic reconstructed voxel size of 251 μm. All PET images were also corrected for random counts, dead time and decay. Images were rendered using Vivoquant v3.0 software (Invicro, USA). Analysis was performed as previously described. Briefly, volumes Of Interest (VOIs) were drawn on the CT images to encompass the brain. A spherical VOI was also outlined on the leg muscle to get the quantification of circulating body radioactivity within the animal. The Mean was extracted from those two VOIs, and the associated mean was calculated.

### RNA isolation and RT-qPCR

The dorsal telencephalon of embryos was micro-dissected in ice-cold RNAse-free 1X PBS. RNA was isolated using the RNA extraction kit (Monarch Total RNA Miniprep Kit, New England Biolabs, Cat# T2010S). 500ng of RNA was reverse transcribed using the reverse transcription kit (iScript cDNA synthesis kit, Biorad, Cat# 1708891). The resultant cDNA was diluted, and reactions were prepared using LUNA qPCR Syber Green Master Mix (New England Biolabs, Cat# M3003). GAPDH was used as a control, 2^-ΔΔCT^ method was used to estimate relative abundance of mRNA. Graphs were plotted with Graphpad PRISM(Version 9), and an unpaired two-tailed Student t-test was used to infer statistical significance. Primers are provided in supplementary information. RT-qPCR primers used were as follows: *Wif1* F 5’-GTGAACTCAGCAAATGCCCC -3’, *Wif1* R *5’- CTCTCGACACTGGCACTTGT-3’ (*Sun et al., 2022*), Hey2* F *5’-* AAG CGC CCT TGT GAG GAA A -3’, *Hey2* R *5’-*TCG CTC CCC ACG TCG AT- 3’ (Benito-Gonzalez et al., 2014), *Dkk3* F *5’-* TGAGGCAGTGGCTACACAAG, *Dkk3* R *5’-* GCTGGTATGGGGTTGAGAGA-3’ (Yin et al., 2018), *Gmnc* F 5’- TGGTCTCCTGGACAACACTG- 3’ *Gmnc* R TAACTCAGAGGGCGATTCCA (Omiya et al., 2021), *Prdm12* F 5’- CGGAATGAGCAGGAGCAGAA-3’, *Prdm12* R 5’-CTCCTGGTCTGGAGGGATCA -3’ (Latragna et al., 2022), *Prdm16* F 5’- CACCGGGTCAGAGGAGAAAT, *Prdm16* R 5’- CGACATGTCAGGGCTCCTAT -3’ (He et al., 2021), *Nfib* F 5’- GGGACTAAGCCCAAGAGACC-3’, *Nfib* R 5’- GTCCAGTCACAAATCCTCAGC-3’ (Wu et al., 2016), GAPDH F 5’- CTCCCACTCTTCCACCTTCG -3’, GAPDH R 5’-GCCTCTCTTGCTCAGTGTCC 3’ (Latragna et al., 2022), *Tox* F 5’-GGCGCGCCATGGACGTAAGATTTTAT-3’, *Tox* R 5’- TTCGAACCCGGTGAGATACAGCGCTTT-3’ (This study), *Shisa2* F 5’- GCGACAACGACCGCCA-3’, *Shisa2* R *5’-* TGAGGAACGGCACGTAGATG(This study), *Fut9* F 5’- TCACGCATCTGATAGCACCG-3’, *Fut9* R 5’-GGGCGAAGAATGCCTTTGGA- 3’ (This study), *Dach1* F 5’- ACGCCAGTTCCAGACCTG-3’, *Dach1* R 5- ATTCCAGGAGACATGAGGCC-3’ (This study).

### Bulk RNA-sequencing and data analysis

E12.5 dorsal telencephalon was harvested in ice-cold PBS and stored in Trizol reagent at -80 °C. RNA extraction was done using illustra RNAspin Mini RNA isolation kit (GE Healthcare, Cat# 25-0500-70). RNA quality was assessed using a sensitivity RNA Analysis kit (DNF-471). Bulk RNA-Seq libraries were performed using 100ng of total RNA of a nonstranded RNA Seq massively parallel mRNA sequencing approach from Illumina (TruSeq RNA Library Preparation Kit v2, Set A; 48 samples, 12 indexes, Cat# RS-122-2001). Libraries were prepared on the automation Beckman Coulter’s Biomek FXP workstation. Accurate quantification of cDNA libraries were done using QuantiFluordsDNA System, a fluorometric-based system, from Promega (Madison, WI). CDNA library size was determined by dsDNA 905 Reagent Kit (Fragment Analyzer from Advanced Bioanalytical) and indicated an average bp length of 280. Pooled Libraries were sequenced on Illumina NovaSeq6000 (PE; 50 bp; 30 Mio reads/sample). BaseCaller software(Illumina) was used to transfer sequence images to to BCL files and subsequently demultiplexed to fastq files using bcl2fastq v2.20.0.422. Sequence quality was ensured using FastQC software (http://www.bioinformatics.babraham.ac.uk/projects/ fastqc/)(version 0.11.9). Sequences were then aligned to the reference *Mus musculus* genome (GRCm39 version 107, https://www. ensembl.org/Mus_musculus/Info/Index) with the help of STAR aligner software version 2.7.8a (Dobin et al., 2013), with a cut off two mismatches within 50bps. Thereafter, read counting was performed using featureCounts version 1.6.3(Liao et al., 2014). Read counts were analysed in the R/ Bioconductor environment version 4.2.2 (www.bioconductor.org) using the DESeq2 package version 1.38.2 (Love et al.,2014). Candidate genes were selected as those with an FDR-corrected P-value of 0.05. Genes were annotated using the *M musculus* GTF file GRCm39 version 107, which was used to quantify the reads within genes. gprofiler2 from ClusterProfiler was used for GO analysis (Kolberg et al., 2020). Gene set enrichment analysis (GSEA) was performed as previously described (Subramanian et al., 2005). Volcano plots were created using EnhancedVolcano package(Blighe et al., 2019). Heatmap were plotted using heatmap.2 package using normalised reads (Gregory et al., 2015).

### Chromatin Immunoprecipitation and ChIP-qPCR<colcnt=5>

E12.5 dorsal telencephalon was dissected in ice-cold PBS. The tissue was fixed for 15 minutes in 1% methanol-free formaldehyde (Thermoscientifc, Cat# 28906) in 1X PBS at room temperature. The crosslinking reaction was stopped by adding Glycine to a final concentration of 125mM and incubated for 8 minutes. The tissue was washed two times before either storing at -80°C or immediately lysed in SDS lysis buffer (50mM Tris Tis pH 8.1, 10mM EDTA, 1% SDS, cOmplete EDTA-free Protease Inhibitor Cocktail, Roche, Cat# 05892791001) on ice for 15 minutes and subjected to sonication for 4 separate cycles of 30s on and 30s off cycles at high setting (Bioruptor Plus, Diagenode). The resultant chromatin was centrifuged at 12000 rpm for 30 minutes to remove any debris. The supernatant was collected and diluted in a ChIP dilution buffer (16.7mM Tris pH8.1, 2mM EDTA, 0.1% SDS (wt/vol), 1% Triton X100(vol/vol), 167mM NaCl, cOmplete EDTA-free Protease Inhibitor Cocktail, Roche, Cat# 05892791001). 10% of the input was de-crosslinked in ten times the volume of input, in elution buffer (1% SDS (wt/vol), 100mM NaHCO3, 200mM NaCl) by incubating at 65°C overnight. The de-crosslinked DNA was purified using the extraction kit (Roche, Cat# 11732676001). 10% of the purified input was run on an agarose gel to determine the size of the chromatin; once a chromatin size of 200-400 bp was confirmed, proceed to the next step. For Emx2 ChIP 4μg of total chromatin and for Ldb1 ChIP 8μg of chromatin was used. Required chromatin was diluted in 4 times more ChIP dilution buffer and incubated with 5μg antibody overnight at 4°C. The next day 30ul ChIP grade Protein G magnetic beads (Cell signalling, Cat# 9906) were added and incubated for 2-3hours before washing two times with low salt buffer (20mM Tris pH8.1, 2mM EDTA, 1% SDS (wt/vol), 200mM) and on time in high salt buffer (20mM Tris pH8.1, 2mM EDTA, 1% SDS (wt/vol), 0.5M NaCl). The bound chromatin was subsequently eluted in elution buffer without NaCl by vigorous shaking at 65°C for 30 minutes. Eluted chromatin was de-crosslinked as described above. The DNA was subsequently purified using kit (Roche, Cat# 11732676001). For each primer pair, relative quantification using the standard curve method was used on a StepOnePlus PCR instrument (Applied Biosystem, Cat# 4376600). Each Primer efficiency was precalculated. Fold enrichment was calculated as the percentage of input values. ChIP-qPCR primers used were as follows: *Sox2* 5’enhancer F 5’-TGGCGAGTGGTTAAACAGAG-3’, *Sox2* 5’enhancer R 5’-CTTATGGAAATGAAGGCGAACG-3’, *Sox2* 3’enhancer F 5’-GCAGCCATTGTGATGCATATAG-3’, *Sox2* 3’enhancer R 5’-CCCGTCATTTGGGTCTTTATT-3’, *Dach1* enhancer F 5’- TGTCTGCATTCTGTCCCTGA -3’, *Dach1* enhancer R 5’- CACCCTTTGATACAGACACTCAC-3’, *Dach1* promoter F 5’- GGTACAAACAATGCCCCAGG-3’, *Dach1* promoter R 5’- TGGCCATAGAAGTGTGCTCA-3’, *Dach1* neg F 5’- AGAAGCAAGTTGGGGAGGAA-3’, *Dach1* neg R 5’- TGACCTGCTTGAGTTCCAGT-3’, *Nfib* enhancer F 5’- TCAGACCTTGTGTGCACTGT-3’, *Nfib* enhancer R 5’- GAAGACCTTGGGATGCCAGT-3’, *Nfib* promoter F 5’- TAACCCCGTCCTGTCCTCAT-3’, *Nfib* promoter R 5’- TGGTAGACTTGGCAAGCTCG, *Nfib* neg F 5’- TAACCCCGTCCTGTCCTCAT-3’, *Nfib* neg R 5’- AGCAGATGGTTTAGGTCGGT, *Prdm16* enhancer F 5’- ATCAGATAACTGCCCCGCTC -3’, *Prdm16* enhancer R 5’- GGGAGCAATTAACAGGCTGC-3’, *Prdm16* neg F 5’- AATAATCGCTGCCCAAAGCC-3’, *Prdm16* neg R 5’- CCTCATGGCTCACGAAGGAT-3’, *Tox* enhancer F 5’- GCCAACTCATCTTCCAAGCC-3’, *Tox* enhancer R 5’- TGTGTGAGGGATTCTGCCAT- 3’, *Tox* neg F 5’- TAAGGTACCAGACCCCGCTA-3’, *Tox* neg R 5’- GGCTAGCCTCCACCTATCCT-3’.

### ChIP-Seq data analysis

Paired-end reads were trimmed using TrimmomaticPE to remove Illumina universal adapters. Trimmed reads were mapped to mouse genome mm10 using Bowtie2 with default parameters for paired-end sequencing (Langmead & Salzberg, 2012). Duplicate reads were removed with MarkDuplicates tools (Picardsuite). Peaks were called with the callpeak tool from MACS2 package with following parameters: -f BAMPE -g mm -q 0.05 –nomodel –call-summits -B –SPMR (Zhang etal., 2008). HOMER’s annotatePeaks.pl tool was used to identify genomic location and gene type for each peak with default parameters. For motifs enrichment analysis, the MEME Suite (MEME-ChIP) (Bailey et al., 2015) and HOMER package were used with default parameters for motif’s length of 8,10,12 bp. For longer motif length, HOMER parameter was changed to 16,18,20 bp (Heinz et al., 2010). BETA package with default parameters was used to infer a regulatory potential of EMX2 and DMRTA5 on the expression of genes annotated to their binding sites (Wang et al., 2013). For Gene ontology analysis, we introduced BED files of Emx2 or Dmrta2 peaks to the GREAT tool with default parameters using Basal plus extension argument with the following parameters: Proximal: 5.0 kb upstream, 1.0 kb downstream, plus Distal: up to 300.0 kb (McLean et al., 2010). For visualisation, deeptools package was used to generate bigwig files and heatmaps (Ramírez et al., 2014). The following publicly available histone marks ChIP-Seq data has been used in this study: GSE82761, GSE83054, GSE82528.

### Immunoprecipitation

P19 cells were transiently transfected Emx2-3x-Flag-V5 construct. 36 hours after transfection, cells were harvested and washed 2 times in ice-cold PBS and lysed using Lysis buffer (150 mM NaCl, 5 mM EDTA pH 8.0, 50 mM Tris pH8.0, 1% NP-40 (vol/vol) supplemented with protease inhibitor cocktail (cOmplete EDTA-free Protease Inhibitor Cocktail, Roche, Cat# 05892791001) for 20 minutes at 4°C. The lysate was then centrifuged at 13000rpm for 30 minutes to remove debris, and the supernatant was used to estimate protein concentration (Biorad, Cat# 5000111). The lysate was checked for expression of tagged Emx2 construct. 4 mg of total protein was used for each experiment. 10 μg of V5 antibody or Normal Rabbit IgG were used per sample and incubated overnight at 4°C. The next day, 100 μl of Protein G magnetic beads (Cell signalling, Cat# 9906) was added and incubated for another 3 hours. Beads were then given a stringent wash of five times using lysis buffer and subsequently sent for mass spectrometry analysis. Cortical lysates were made using the process mentioned above. 2mg of total protein was used for immune precipitation, and 3μg of antibody was used per IP. Following the above-mentioned process of immunoprecipitation, beads were subjected to boiling in Laemmli.sample buffer at 90°C for 10 minutes. Eluted proteins were subsequently subjected to immunoblotting.

For co-immuno precipitation, transfected HEK293T cells (CalPhos Mammalian Transfection Kit, Cat# 631312, Takara) were harvested 236 hours after transfection. Lysis buffer (150 mM NaCl, 5 mM EDTA pH 8.0, 50 mM Tris pH8.0, 1% NP-40) supplemented with protease inhibitor cocktail (cOmplete EDTA-free Protease Inhibitor Cocktail, Roche, Cat# 05892791001) was added, and cells were left to incubate for 15 minutes at 4°C. The collected cell lysate was centrifuged for 20 minutes at 13000rpm. Co-IPs were performed by incubating cell lysate(1mg) with the required antibody (3-5 μg) for overnight at 4°C. Protein G magnetic beads (Cell signalling, Cat# 9906) were added the next day and incubated for at least 2-3 hours before washing five times with lysis buffer supplemented with protease inhibitor cocktail. Immunoprecipitated proteins were eluted by incubating beads in Laemmli sample buffer at 90°C for 10 minutes. Eluted proteins were subsequently subjected to immunoblotting.

### Rapid immunoprecipitation mass spectrometry of endogenous proteins

Rapid immunoprecipitation mass spectrometry of endogenous proteins (RIME) was carried out as previously described with minor modifications (Mohammed et al., 2016). E12.5 cortices were dissected on ice-cold PBS and fixed for 15 minutes in 1% methanol-free formaldehyde (Thermoscientifc, Cat# 28906) in PBS at RT. The cross-linking was quenched by adding glycine to a final concentration of 125mM and incubating samples for 8 minutes. Subsequently, cells were pelleted and washed twice in ice-cold PBS. Nuclear extraction was done using ice-cold LB1 buffer (150mM HEPES-KOH pH7.5, 140mM NaCl, 1mM EDTA, 10% glycerol, 0.5% NP-40 (vol/vol) and 0.25% Triton X-100(vol/vol)), incubating cells on rotation at 4°C for 10 minutes. The suspension was then centrifuged at 2000g for 5 minutes at 4°C, and the obtained nuclear pellet was washed in ice-cold LB2 buffer (10 mM Tris-HCl pH 8.0, 200 mM NaCl, 1 mM EDTA, 0.5 mM EGTA). Pelleted nuclei were subsequently resuspended in LB3 buffer (10 mM Tris-HCl pH 8.0, 100 mM NaCl, 1mM EDTA, 0.5 mM EGTA, 0.1% sodium deoxycholate (wt/vol) and 0.5% N-lauroylsarcosine (wt/vol)) and, after a 15 minutes incubation, were subsequently subjected to 3 cycles of sonication (30s ON/30s OFF for 10 minutes at 200W, high setting, Bioruptor, Diagenode). Triton X-100 was added to a final concentration of 1%. The obtained chromatin was then cleared by centrifuging at 20.000g for 4 minutes and the supernatant was collected. The obtained chromatin was de-crosslinked overnight to assess the size of the fragments. 10% of the input was de-crosslinked in ten times the volume of input, in elution buffer (1% SDS, 100mM NaHCO_3_, 200mM NaCl) by incubating at 65°C overnight. The de-crosslinked DNA was purified using the extraction kit (Roche, Cat# 11732676001). DNA was checked on an agarose gel to see 200-600bp-sized fragments. Antibody-bound beads were generated by incubating 100μl of Protein G magnetic beads (Cell signalling, Cat# 9906) with 10μg anti-V5 antibody (Abcam, Cat# ab15828) overnight at 4°C. The next day, the antibody-bound beads were washed three times to remove any unbound antibodies. For immunoprecipitation, 2mg of cortical protein lysates were used per RIME and incubated overnight at 4°Cin RIPA buffer (50 mM HEPES pH 7.6, 1 mM EDTA, 0.7% (wt/vol) sodium deoxycholate, 1% (vol/vol) NP-40 and 0.5M LiCl). Subsequently, the chromatin-bound beads were washed in RIPA buffer for 9 washes and then washed two times more with 100 mM ammonium hydrogen carbonate (AMBIC) solution. Beads were stored at -80°C until Mass spectrometry analysis.

### Processing of RIME and IP-MS samples by Mass spectroscopy

Protein digestion was performed in the following way: 100 µl of formic acid 1 % were placed in each Millipore Microcon 30 MRCFOR030 Ultracel PL-30 to wash the filter and centrifuged for 15 min at 14500 rpm. 40 µg of protein adjusted in 150 µl of urea buffer 8M (urea 8 M in buffer Tris 0.1 M at pH 8,5) were placed individually in a column corresponding to each sample and centrifuged for 15 min at 14500 rpm. After discarding the filtrate, 200 µl of urea buffer was added to the columns and washed three times by centrifugation at 14500 rpm for 15 minutes. For the reduction, 100 µl of dithiothreitol (DTT) were added to the column and mixed for a minute in a thermomixer and subsequently incubated for 15 minutes at 24 °C. Then the samples were centrifugated at 14500 rpm for 15 minutes and the filtrate was discarded. 100 µl of urea buffer was added to clean the filter by centrifuging at 14500 rpm for 15 minutes. Alkylation was performed by adding 100 µl of iodoacetamide (IAA), in urea buffer) in the column and mixing at 400 rpm for 1 minute in the dark and which column was incubated in the dark for 20 minutes and centrifuged for 10 minutes at 14500 rpm. Excess IAA was removed by adding 100 µl of urea buffer, and samples were centrifuged for 15 minutes at 14500 rpm. To quench the residual IAA, 100 µl of DTT was added to the column, mixed for a minute at 400 rpm, and incubated at 24 °C for 15 minutes and then centrifuged for 10 minutes at 14500 rpm. The filtrate was discarded, the column was washed thrice by adding 100 µl of sodium bicarbonate buffer 50 mM ((ABC) in ultrapure water), then centrifuging for 10 minutes at 14500 rpm. In the last round, the remaining 100 µl was kept at the bottom of the column to avoid evaporation in the column. Peptide digestion was done by adding 80 µl of mass spectrometry-grade trypsin (1/50 in ABC buffer) to the column, mixing for a minute at 400 rpm, and then incubating the sample at 24°C in a water-saturated environment overnight. The next day, the columns were placed in a 1.5 ml LoBind tube and centrifuged for 10 minutes at 14500 rpm before adding 40 µl of ABC buffer to the column. 10 % Trifluoroacetic acid (TFA) in ultrapure water was added to the extract of the LoBind Tube to obtain 0.2 % TFA. The samples were dried in a SpeedVac up to 20 µl and transferred in an injection vial.

The digested samples were analyzed using nano-LC-ESI-MS/MS tims TOF Pro (Bruker, Billerica, MA, USA) coupled with an UHPLC nanoElute (Bruker). Peptides were separated by nanoUHPLC (nanoElute, Bruker) on a 75 μm ID, 25 cm C18 column with integrated CaptiveSpray insert (Aurora, ionopticks, Melbourne) at a flow rate of 200 nl/min, at 50°C. LC mobile phases A were water with 0.1% formic acid (v/v) and B ACN with formic acid 0.1% (v/v). Samples were loaded directly on the analytical column at a constant pressure of 800 bar. The digest (1 µl) was injected, and the organic content of the mobile phase was increased linearly from 2% B to 15 % in 40 min, from 15 % B to 25% in 15 min, from 25% B to 37 % in 10 min and from 37% B to 95% in 5 min. Data acquisition on the tims TOF Pro was performed using Hystar 6.1 and timsControl 2.0. tims TOF Pro data were acquired using 160 ms TIMS accumulation time, mobility (1/K0) range from 0.75 to 1.42 Vs/cm². Mass-spectrometry analysis was carried out using the parallel accumulation serial fragmentation (PASEF) (Meier et al., 2018) acquisition method. One MS spectra followed by PASEF MSMS spectra per total cycle of 1.16 s.

Data analysis was performed using Mascot 2.8.1 (Matrix Science). Tandem mass spectra were extracted, charge state deconvoluted and deisotoped by Data analysis (Bruker) version 5.3. All MS/MS samples were analyzed using Mascot (Matrix Science, London, UK; version 2.8.1). Mascot was set up to search the *Mus musculus* database from Uniprot (97856 entries) assuming the digestion enzyme trypsin. Mascot was searched with a fragment ion mass tolerance of 0.050 Da and a parent ion tolerance of 15 PPM. Carbamidomethyl of cysteine was specified in Mascot as fixed modifications. Oxidation of methionine and acetyl of the n-terminus were specified in Mascot as variable modifications. Scaffold (version Scaffold_5.0.0, Proteome Software Inc., Portland, OR) was used to validate MS/MS-based peptide and protein identifications. Peptide identifications were accepted if they could be established at greater than 96.0% probability to achieve an FDR less than 1.0% by the Scaffold Local FDR algorithm. Protein identifications were accepted if they could be established at greater than 5.0% probability to achieve an FDR less than 1.0% and contained at least 2 identified peptides. Protein probabilities were assigned by the Protein Prophet algorithm (Nesvizhskii et al., 2003). Proteins that contained similar peptides and could not be differentiated based on MS/MS analysis alone were grouped to satisfy the principles of parsimony. Proteins sharing significant peptide evidence were grouped into clusters. Volcano plots were created using EnhancedVolcano package (Blighe et al., 2019). Heatmap were plotted using heatmap.2 package using a normalised total number of spectra (Warnes et al., 2020).

### SEM and TEM analysis

Immediately after dissection, brains were fixed overnight at room temperature (RT) in a 0.1M cacodylate buffer solution containing 2% paraformaldehyde and 2.5% glutaraldehyde. Hemispherectomy was performed, and one hemisphere was processed for scanning electron microscopy (SEM) while the other half was prepared for transmission electron microscopy (TEM). For SEM, hemispheres were additionally fixed in 2.5% glutaraldehyde in cacodylate buffer at RT for a few hours, then immobilized in 2% low melting point agarose and sectioned using a vibratome. Sections of 150 to 300 µm thickness were obtained, fixed in 2.5% glutaraldehyde for an additional 2 hours at RT, and post-fixed in 2% osmium tetroxide in cacodylate buffer for 1 hour at RT. Samples were then gradually dehydrated in a series of acetone baths (ranging from 50% to 100%, 15 minutes each) and dried using critical point drying (CPD 030, Leica). Once mounted on SEM supports, samples were coated with a thin layer of platinum (20 nm, MED020, Leica) and examined using a Quanta F200 SEM microscope (FEI) at 30 kV with a secondary electron detector.

For TEM, hemispheres were dissected and fixed in 2.5% glutaraldehyde for a few (1 to 3) hours several times to encompass the minimal region surrounding the choroid plexus. For KO mice, the cortex was removed, exposing the region of interest. For WT mice, consecutive regions exposing the lateral ventricles were sectioned and processed in parallel. Samples were then washed in cacodylate buffer, post-fixed in 1% osmium tetroxide and 1.5% potassium ferrocyanide for 1 hour at RT, washed again, then immersed in an additional bath of 1% osmium tetroxide for 1 hour, rinsed in water, and stained with 1% uranyl acetate solution for 2 hours at RT. After washing, samples were dehydrated in a series of ethanol solutions (ranging from 35% to 100%, 15 minutes each). Ethanol was replaced with 100% propylene oxide, followed by infiltration with epoxy resin (AGAR100, Agar Scientific). Once embedded in resin, samples were placed in an oven at 60°C for 72 hours. Sections were then obtained using a UC6 ultramicrotome (Leica). Semi-thin sections were initially prepared to confirm localization in the choroid plexus, followed by ultra-thin sections (50 to 70 nm) mounted on copper grids (100 mesh, formvar/carbon-coated). Grids were examined using a Tecnai 10 100 kV microscope (FEI) equipped with a Veleta 14-bit camera.

### Statistical Analysis

All statistical tests were performed in GraphPad version 9. Appropriate statistical tests were chosen based on sample size and possible sample distribution. No statistical test was performed to infer sample distribution or its nature, and no test was performed to pre-determine the sample size. The chosen statistical test and the p-values obtained are mentioned in respective legends.

## RESULTS

### *Dmrta2* and *Emx2* regulate a similar set of genes in the dorsal telencephalon

*Dmrta2* and *Emx2* are expressed in a high caudo-medial to low rostro-lateral gradient along the ventricular zone in the developing cortex (**Fig. 1A**). Multiple double mutants have been generated previously to understand the cooperation between *Emx2* and other signaling molecule or TFs during cortex development, including *Emx1, Pax6, Otx2, Otx1, Fgf17, Foxg1, Dmrta2* (Shinozaki et al., 2002, Bishop et al. 2002, Muzio et al., 2002a, b; Muzio & Mallamaci, 2003; Shinozaki et al., 2004; Muzio et al., 2005; Kimura et al., 2005; Cholfin et al., 2008; Desmaris et al., 2018). Among them, the combinatorial deletion of *Emx2* with *Pax6* or *Dmrta2* has been shown to impart a much adverse phenotype regarding cortical size. However, in the case of *Emx2;Dmrta2* double mutants, our previous analysis was limited to E12.5 embryos (Desmaris et al., 2018). To further evaluate the extent of the cortical phenotype of *Emx2;Dmrta2* double mutants, we analysed their cortical size at E18.5. As previously described, we observed that *Emx2* single (33.82 ± 3.94 %, n = 8, 7; p<0.0001) mutants and *Dmrta2* single mutants (51.70 ± 4.00 %, n = 8, 5; p<0.0001) have reduced cortices compared to WT, the reduction being more severe in *Dmrta2* than in *Emx2* single mutants (27.81 ± 6.92 %, n= 7, 5; p = 0.0025). In *Emx2*^-/-^;*Dmrta2^+/-^* mutants, the cortex was smaller than in *Emx2* single mutants (22.42 ± 6.93%, n= 7, 5, p=0.0090). In contrast, *Emx2*^+/-^; *Dmrta2^-/-^* mutants did not show a more severe reduction in the size of their cortex than *Dmrta2* single mutants. Notably, the combined loss of *Dmrta2* and *Emx2* led to a complete loss of the cortex (100%, n=4, p<0.0001) (**Fig. 1B, C**). These results further suggest that *Emx2* and *Dmrta2* function together to regulate cortical development.

**Figure 1:**
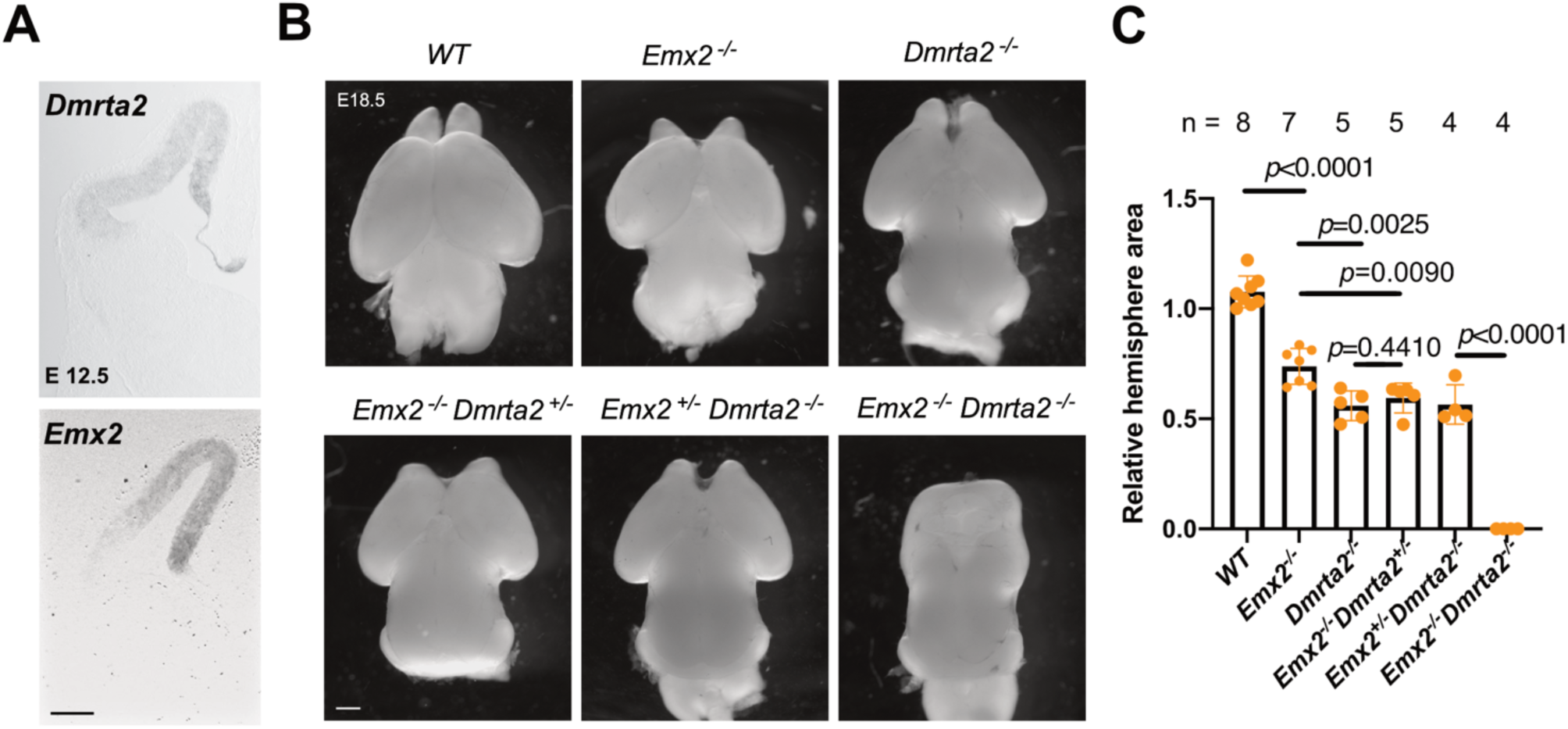
Ablation of both *Dmrta2* and *Emx2* leads to a complete loss of the cortex at E18.5. **A**. Section of the telencephalon of E12.5 embryos at a caudal level, showing by ISH *Dmrta2* and *Emx2* medial high lateral low expression gradients(scale bar 100μm). **B**. Dorsal views of the brain of E18.5 embryos of the indicated genotypes (scale bar 100μm). **C**. Relative dorsal hemisphere size of embryos with the indicated genotypes. Error bars showing mean ± SD. Each point on the graph represents an independent biological replicate. (genotype, n, mean ± SD) (*WT*, 8, 1.076± 0.071), (*Emx2*^-/-^, 7, 0.738 ±0.081), (*Dmrta2^-/-^*, 5, 0.559 ± 0.067), (*Emx2^-/-^Dmrta2^+/-^*, 5, 0.593 ± 0.067), (*Emx2^+/-^Dmrta2^-/-^*, 4, 0.564 ± 0.088), (*Emx2^-/-^Dmrta2^-/-^*, 4). Unpaired two-tailed T-test were used to calculate *p* values for all comparisons except for *Emx2^+/-^Dmrta2^-/-^*vs *Emx2^-/-^Dmrta2^-/-^* for which unpaired two-tailed T-test with Welch’s correction was used.

Though *Emx2* is one of the most studied TF during cortical development (Pellegrini et al., 1996; Yoshida et al., 1997; Rhodes et al., 2003; Kimura et al., 2005; Dwyer et al., 2011), a comprehensive understanding of deregulated genes in *Emx2* KO is lacking. Therefore, we analyzed by bulk RNA-seq the transcriptome of dorsal telencephalic tissue dissected from E12.5 *Emx2* knockout embryos. As expected, transcripts corresponding to the end of exon 2 were absent in *Emx2* mutants (Pellegrini et al., 1996) (**Extended Data** Figure 1-1A). In these *Emx2* mutants, we found that many genes known to be negatively regulated by *Emx2*, including *Pax6*, are only marginally upregulated in terms of fold change (FC = 1.25; padj= 0.0001), likely due to their gradient expression within the telencephalon, wherein a slight change in dosage is sufficient to shift the gradients to impart a phenotype evident from the arealisation defect in *Emx2* heterozygous mice (Leingärtner et al., 2007). To confirm their deregulation, we analysed by qRT-PCR the expression of several genes expressed in gradient across the telencephalon, including *Tox* (lateral and ventral pallium, subpallium), *Prdm12* (pallial-subpallial boundary), *Shisa2* (dorsal pallium), *Wif1*, *Fut9*, *Hey2* (medial pallium), *Dkk3* (cortical hem) and *Gmnc* (cortical hem, choroid plexus and Cajal-Retzius cells) (Leimeister et al., 1999; Kinameri et al., 2008; Artegiani et al., 2015; Abdullah et al., 2022; Moreau et al., 2023). *Tox*, *Prdm12* and *Shisa2* are expressed in an opposed gradient to that of *Emx2* and were upregulated in *Emx2* KO, whereas *Wif1*, *Fut9*, *Hey2*, *Dkk3* and *Gmnc* that are expressed in a similar gradient as *Emx2* were downregulated in *Emx2* KO (**Extended Data** Figure 1-1B).

We subsequently compared the bulk RNA seq data obtained from cortices of *Emx2* KO embryos with that of *Dmrta2* KO embryos (Desmaris et al., 2018). Among the top 100 downregulated genes, 42 genes were shared between *Dmrta2* KO and *Emx2* KO. Among them are genes that are expressed in the cortical hem and Cajal-Retzius cells (*Wnt2b*, *Wnt3a*, *Rspo1*) and choroid plexus (*Ttr*, *Ccno*, *Gmnc*, *Sulf1*, *Mcidas*). This is consistent with *Emx2* and *Dmrta2*’s role in maintaining Wnt-rich cortical hem structure and in the development of Cajal-Retzius cells (Shinozaki et al., 2002; Shinozaki et al., 2004; Saulnier et al., 2013). On the other hand, of the top 100 most upregulated genes, 18 were common between *Dmrta2* KO and *Emx2* KO. The common upregulated genes are expressed in the lateral and ventral pallium and subpallium, with subpallial genes exhibiting a higher fold change. The genes upregulated include *Tox* (lateral and ventral pallium subpallium), *Prdm12*, *Fgf3* (pallial-subpallial boundary), *Pou3f1* (lateral and dorsal pallium), *Gsx2*, *Meis1*, *Isl1*, *Prdm13* (subpallium), *Neurog1*(lateral pallium and ventral pallium) (**Fig. 2 A-C, E)**. Gene Ontology (GO) analysis indicated that similar biological processes are deregulated in both single knockouts (**Fig. 2 D, F**). The most downregulated genes in both cases belong to genes related to cilium organisation, cilium-dependent cell mobility, cilium assembly, and cilium movement. This corresponds to genes expressed in the choroid plexus (Terré et al., 2016) and the cortical hem-derived Cajal-Retzius cells that transiently express multiciliated genes (Moreau et al., 2023). GO terms for genes upregulated in both knockouts were related to neurogenesis and nervous system development, neuron differentiation and forebrain development. This is also consistent with the fact that neurogenesis occurs earlier in the lateral pallium compared to the medial pallium (Saito et al., 2019). To gain a more quantitative understanding of gene sets that are deregulated in each bulk RNA data set, Gene Set Enrichment Analysis (GSEA) was performed. This analysis confirmed the enrichment of pathways related to motile cilium assembly and cilium and flagellum-dependant cell mobility in the set of genes identifed as down-regulated in both *Dmrta2* and *Emx2* KO (**Extended Data** Figure 2-1A**, B**). With this analysis, we however have not been able to obtain gene sets that are upregulated in both *Dmrta2* and *Emx2* KO data.

**Figure 2:**
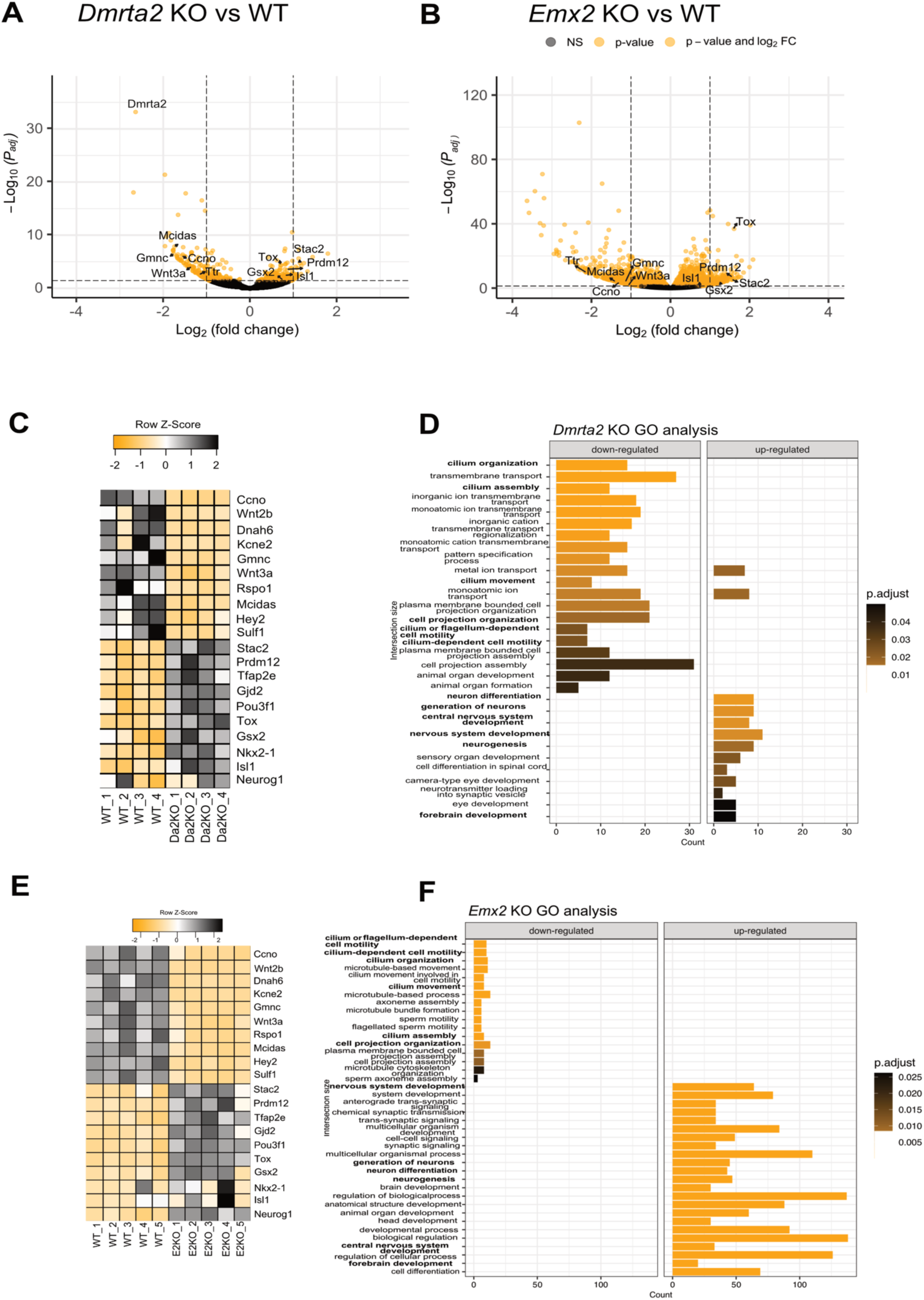
*Dmrta2* and *Emx2* regulate a similar set of genes. **A, B**. Volcano plots showing differentially expressed genes in the respective single knockouts. A *Padj* cut-off of 0.05 and a Log_2_(Fold Change) cut-off ± 1 are drawn by dashed lines. All genes showing significant *Padj* values, irrespective of fold change, are represented as orange points. Otherwise, they are indicated as black points. Five common deregulated genes in the top 100 genes in each data set are indicated with their names. **C, E**. 10 downregulated and upregulated genes in the top 100 genes in both single knockout and their normalised reads are depicted as a heatmap. **D, F**. GO analysis of the differently expressed genes identified using a Log_2_(Fold Change) cut-off ± 1 in indicated single knockouts. The most significantly enriched biological processes are shown. GO terms present in both data sets are highlighted in bold fonts.

We next validated by ISH the deregulation of the expression of some of the identified genes in the telencephalon of E12.5 single and double knockout. This includes genes that are expressed in the medial pallium such as *Gmnc*, *Wnt3a*, *Wnt8b*, and in the vLGE *Isl1* (**Extended Data** Figure 2-2). Results obtained confirmed the reduction of their expression in the telencephalon of both single knockouts and revealed that they are absent in the double knockout embryos. On the other hand, *Isl1* was restricted to the subpallium in single knockouts but was upregulated in the pallium of the double knockout embryos. Thus, a similar set of genes is deregulated in the dorsal telencephalon of *Dmrta2* and *Emx2* single mutants and the compound loss of *Emx2* and *Dmrta2* leads to a more severe deregulation of their expression.

### Dmrta2 and Emx2 common direct targets are limited but include key regulators of cortical development

The cooperative function of Emx2 and Dmrta2 in the developing telencephalon may be due to direct regulation of some of their common targets. Given that Chromatin Immunoprecipitation followed by Sequencing (ChIP-Seq) data in the developing cortex were available for Dmrta2 (Konno et al., 2019) but not for Emx2 at the time we performed this study, we wanted to identify genome-wide Emx2 direct targets in the cortex using ChIP-Seq. Since ChIP-seq graded Emx2 antibodies were not available to us, we generated an *Emx2* knockin (Emx2 KI) line that faithfully expresses a C-terminal 3xFlag-V5 tagged version of the Emx2 protein (**Fig. 3A** and **Extended Data** Figure 3-1 **A-C**). Emx2 has previously been shown to bind and negatively regulate the activity of 3’ and 5’ *Sox2* enhancers in the telencephalon (Mariani et al., 2012). We used this information to optimise the Emx2 ChIP using V5 antibodies on chromatin prepared from dissected cortices of E12.5 KI and WT controls. ChIP-qPCR performed on *Sox2* 3’and 5’ enhancers on Emx2 KI cortices showed selective enrichment when using V5 antibodies (**Extended Data** Figure 3-1 **D**). No such enrichment was observed using IgG, or using V5 antibodies on WT controls. In subsequent ChIP-Seq experiments, we identified a total of 9478 peaks. Our duplicate showed consistency in called peaks (**Fig. 3B**). Most peaks were intergenic (55.9%), followed by intronic (39.4%). Among gene types, most of the peaks were near protein-coding (73.2%) or long non-coding RNA genes (**Extended Data** Figure 3-1 **E**). *De novo* motif analysis identified a motif containing a TAATTA site well in alignment with the Emx2 motif determined using *in vitro* HT-SELEX sequencing (**Fig. 3C**) (https://jaspar.elixir.no/matrix/MA0886.2/). These results aligned well with a recent independent Emx2 ChIP-Seq study using anti-Emx2 antibody (Ypsilanti et al., 2021). The most commonly known motifs enriched in Emx2 peaks included motifs of Emx1 and Lhx2, which are already known to have a similar expression and related functions in the developing cortex (**Extended Data** Figure 3-1 **F**). Recently, Gsx2, another NKL homeobox protein similar to Emx2, expressed in the developing telencephalon, has been shown to bind as a monomer or homodimer to DNA, contributing to downstream targets’ activation or repression, respectively (Salomone et al., 2021). Similarly, Emx2 has been shown to bind to DNA as a monomer or homodimer via HT-SELEX (Jolma et al., 2013). We thus used HOMER *de novo* and known motif discovery to assess long motifs that might contribute to a similar function for Emx2 but didn’t obtain a long or short motif that can be interpreted as dimeric versus monomeric binding (data not shown).

**Figure 3:**
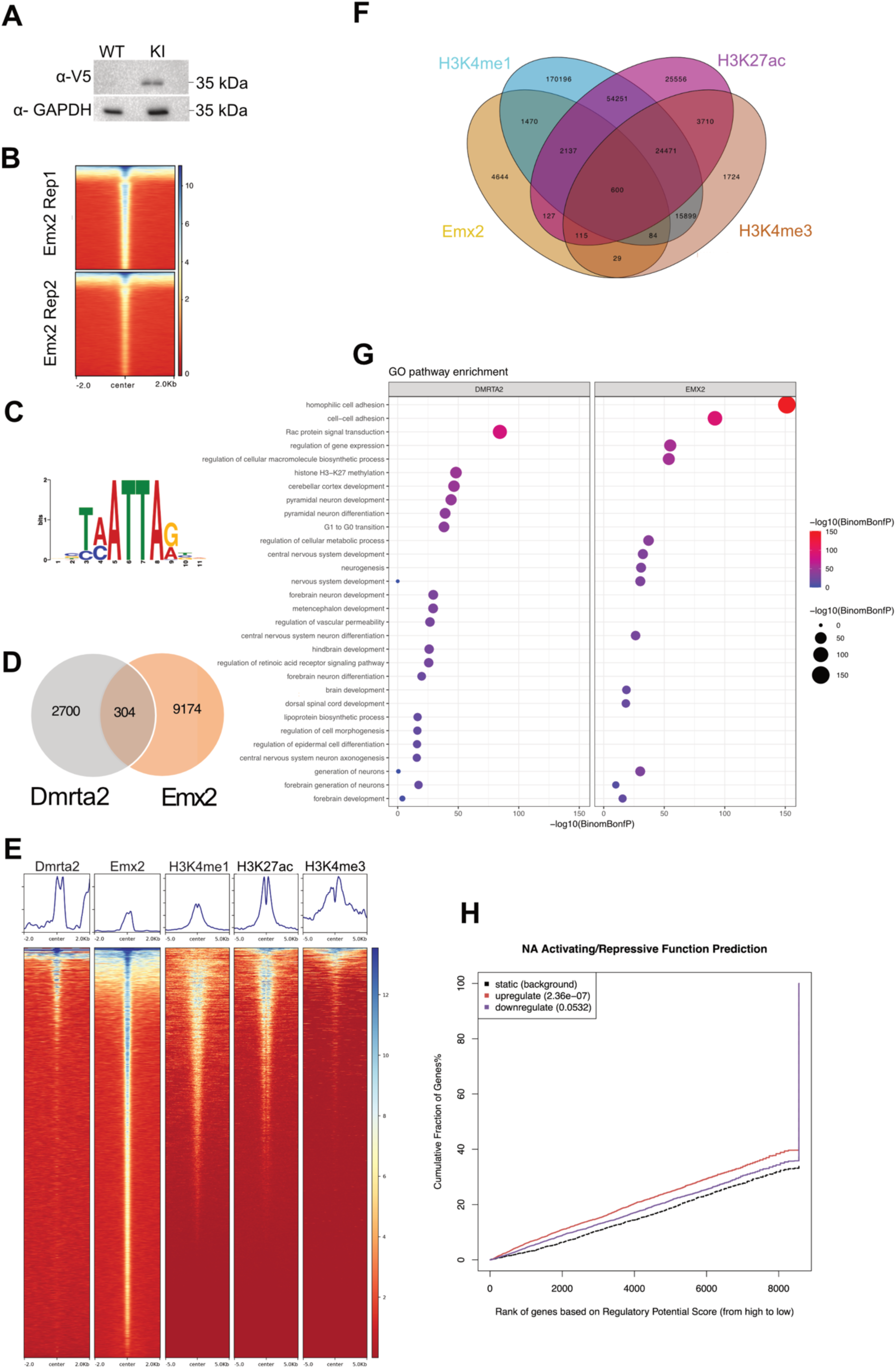
Dmrta2 and Emx2 have a limited number of common direct targets. **A**. Western blot analysis using V5 antibodies showing a signal in a protein extract prepared from dissected cortices from *Emx2 3X FLAG-V5* KI but not from control WT E12.5 embryos. **B**. A heat map comparing the binding pattern of Emx2 across identified peaks in two high-throughput experiments showing consistency between the replicates**. C**. *De novo* motif of Emx2 predicted using MEME from the ChIP-Seq data. **D**. A Venn diagram indicating the shared number of peaks between Emx2 and Dmrta2. **E, F**. Heat maps and a Venn diagram showing the very little overlap of peaks between Dmrta2 and Emx2 and between Emx2 and H3K4me1, H3K27ac and H3K4me3 marks. **G**. GO analysis showing that Dmrta2 or Emx2 mostly regulate distinct pathways. **H**. BETA analysis showing that there is a correlation between upregulated genes identified by RNA-seq in *Emx2* KO embryos and those identified as bound by Emx2 in our ChIP-Seq data.

To compare the peaks between Dmrta2 and Emx2, we utilized the previously published data from Dmrta2 ChIP-Seq (Konno et al., 2019). We observed only 304 peaks common between Dmrta2 and Emx2 ChIP-seq data (**Fig. 3D**). Additionally, we also compared Emx2 peaks with publicly available histone mark ChIP-Seq data in the forebrain. Our analysis revealed 46.61% overlap between Emx2 and H3K4me1 marks, 32.35% overlap and H3K27ac marks and 8.9% overlap with H3K4me3 marks (**Fig. 3E, F**). GO pathway enrichment analysis of genes associated with peaks of Emx2 and Dmrta2 indicated they regulate mostly dissimilar pathways, which is in line with the fact that they share only a subset of common targets (**Fig. 3G**). BETA analysis assesses correspondence between deregulated genes in bulk RNA-Seq performed upon loss of a TF factor and its ChIP-Seq peaks near the deregulated genes. We performed the BETA analysis with the *Emx2* KO RNA-Seq data and the Emx2 ChIP-Seq data. In accordance with the recent observation that most known TFs in the dorsal telencephalon bind directly to the lateral pallial genes rather than to medial pallial genes (Ypsilanti et al., 2021), our analysis showed a strong positive correlation between upregulated genes and Emx2 peaks near them (p<0.005). Although we did find direct binding of Emx2 on some of the downregulated genes (i.e *Emx1*, *Nfix*, *Rspo1*, *Gmnc*, *Foxp2*, *Mytl1*, *Myt1*, and *Axin2*), downregulated genes did not show such a correlation (**Fig. 3H**). BETA analysis on *Dmrta2* KO RNA-Seq against Dmrta2 ChIP-Seq peaks yielded also no significant correlation, both for upregulated and downregulated genes (**Extended Data** Fig. 3-1G).

Our data also shows that Emx2 directly binds to many targets known to be essential for dorsal telencephalon patterning, including *Dmrta2*, *Dmrt3*, *Lhx2*, *Sp8*, *Nr2f1*, *Pbx1*, *Pax6*. Surprisingly, except for *Pax6*, none of the aforementioned genes were directly bound by both Emx2 and Dmrta2 at same enhancer. Emx2 and Dmrta2 have been shown previously by EMSA to bind to a downstream enhancer in the *Gsx2* locus (Desmaris et al., 2018). Our ChIP-Seq data confirm these bindings. They revealed that Emx2 and Dmrta2 bind together to several other genes, most of them being not expressed in the medial telencephalon. Among them, genes expressed in cortical progenitors such as *Dach1*(medial and dorsal pallium), *Foxp4* (dorsal pallium) *Tox*, *Prdm16*, *Zfhx4* (lateral pallium), *Rfx3* (ventral pallium) *Nkx2.1*, *Vax1*, *Zfhx3*, *Nkx6.2*, *Lmo1*, *Hmx3* and *Dlk1* (subpallium) and others expressed post mitotically such as *Zeb2* (lateral pallium), *Gbx1* (subpallium) and *Lhx1* (Cajal Retzius cells).

*Pax6*, *Gsx2* and *Nkx2.1* are essential fate determinants of the lateral pallium, lateral and medial ganglionic eminence, respectively (Sussel et al., 1999; Corbin et al., 2000; Toresson et al., 2000a). *Tox* has an expression pattern similar to that of *Pax6* and has been described as a novel key player in corticogenesis (Artegiani et al., 2015). To determine the impact of Dmrta2 and Emx2 direct binding to these essential telencephalic regulators, we examined their expression in single and double knockouts (**Fig. 4 A-D)**. For *Pax6* and *Tox* we observed only a modest upregulation in *Emx2* KO embryos, and a higher upregulation extending into the medial cortex in *Dmrta2* KO embryos. By contrast, *Pax6* and *Tox* were undetectable in the brain of double KO embryos, which is in accordance with the previous observation that *Ngn2*, a direct Pax6 target, is also lost (Desmaris et al., 2018). Whereas in WT embryos and in both single mutants Gsx2 remains restricted to the subpallium, it extended into the pallium in the double KO. Whereas in the WT embryos and in both single mutants *Nkx2.1* expression domain remains restricted to the MGE, it extended to the LGE in the double mutants. No upregulation of *Nkx2.1* was however observed in the dorsal telencephalon, presumably because the upregulated *Gsx2* can repress it (Corbin et al., 2003). *Nkx2.1* upregulation observed in the LGE in the double knockout could be a non-cell autonomous effect. Indeed, an enlargement of LGE and MGE was observed in the double knockout of *Emx1;Emx2* double knockout (A. Mallamaci, personal communication), indicating that the loss of TFs in the dorsal telencephalon can influence LGE and MGE development. Thus Emx2 and Dmrta2 have only a few direct common targets but some of them encode essential regulators of cortical fate. They suggest that their cooperative repressive action is critical to determine the exact positional information of cortical progenitors.

**Figure 4:**
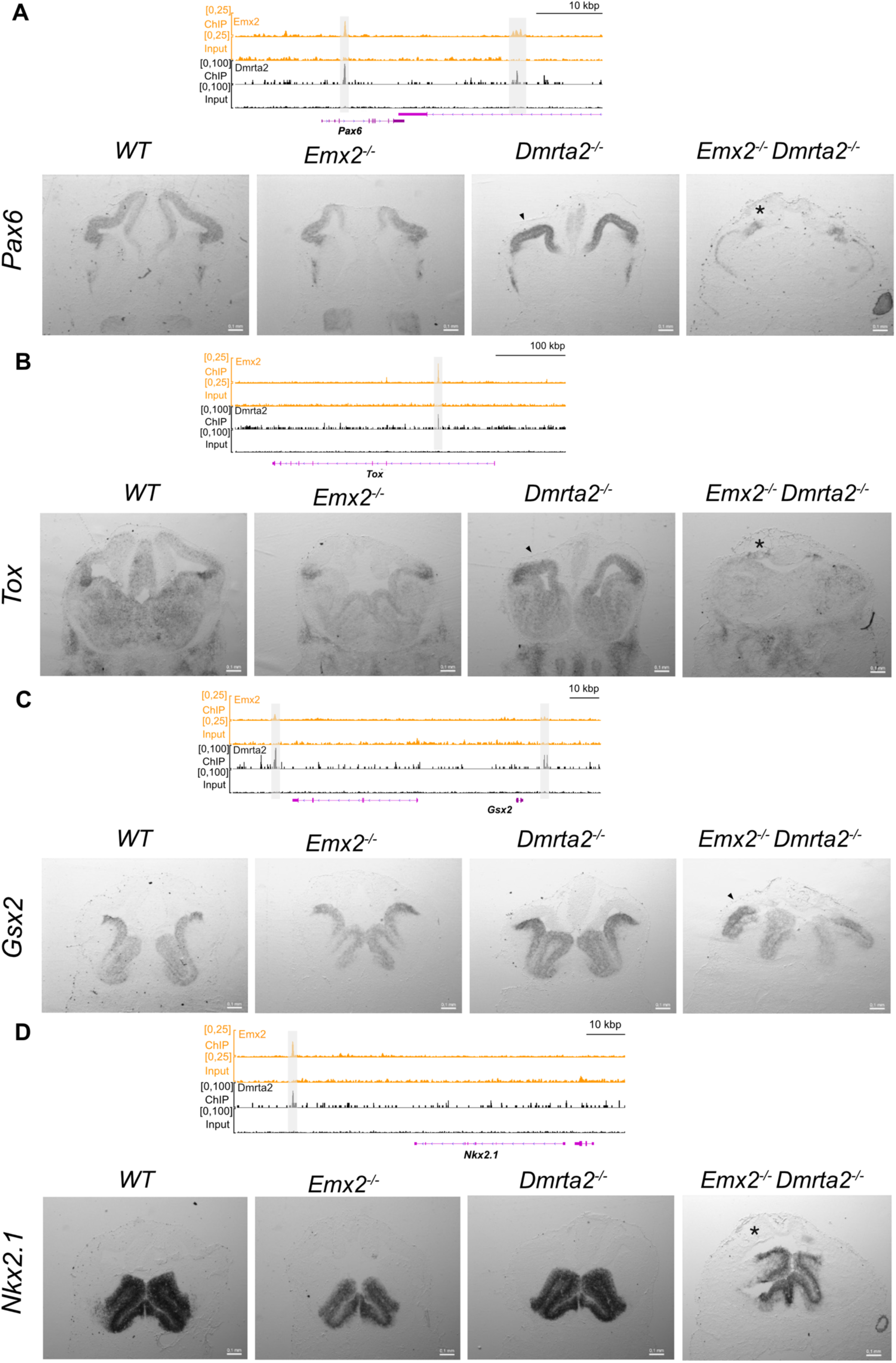
Emx2 and Dmrta2 directly regulate some key cortical determinants. **A-D**. In each panel, IGV snapshots showing Emx2 and Dmrta2 ChIP-Seq peaks around the indicated genes and coronal sections of the cortex of E12.5 single and double KO embryos processed by ISH with antisense probe to examine their expression are shown. Black arrowheads indicate apparent upregulation, and black asterisk indicate near complete loss (scale bar 100μm)(n=3).

### Ldb1 interacts with Emx2 in the developing telencephalon

Currently, no *in vivo* Dmrta2 and Emx2 interacting partners are known during cortical development. Their identification constitutes a key step towards a better understanding of their mechanism of action. As GST pulldown assays have shown that Emx2 can directly bind to the zinc finger transcription factor Sp8 (Zembrzycki et al., 2007), we first tested whether the zinc finger TF Dmrta2 can also interact with Emx2. Results obtained using co-immunoprecipitation assays with overexpressed tagged versions of Emx2 and Dmrta2 produced in HEK293T cells showed they can interact, and that the deletion of its DM domain compromises this interaction (**Fig. 5A**). To identify in an unbiased way Emx2 interactive partners, we then utilized two methods: Rapid immunoprecipitation mass spectrometry of endogenous proteins (RIME) using dissected dorsal telencephalon of Emx2 KI embryos at E12.5 and Immunoprecipitation followed by mass spectrometry (IP-MS) analysis using mouse teratocarcinoma P19 cells transiently overexpressing a 3x-Flag-V5 tagged version of Emx2 (**Fig. 5B**). Using the RIME method on dissected telencephalic tissue from Emx2 KI embryos (**Fig. 5C-D** and **Extended Data Table 5-1**), besides Emx2 peptides, we obtained an enrichment in peptides corresponding to other cortical transcriptional regulators, including Lhx2 and Emx1, expressed in a similar graded manner to Emx2 in cortical progenitors and Pbx1 expressed in an opposite gradient in the ventricular zone of the developing cortex (**Fig. 5E**). Despite the fact that Dmrta2 can interact with Emx2 when overexpressed in cultured cells, we did not observe any Dmrta2 enrichment, which is in accordance with our ChIP-seq data showing that only a limited set of genes are co-bound by Emx2 and Dmrta2. Lhx2 is a likely relevant Emx2 cooperative partner as they have been shown to share a large number of ChIP peaks in the dorsal telencephalon, more than with any other TF (Ypsilanti et al., 2021) and because in our Emx2 ChIP-Seq data, Lhx2 motifs were present in the top 10 motifs. The homologous Emx1 TF is another relevant candidate Emx2 cooperating partner as the compound deletion of *Emx1* and *Emx2* results in adverse phenotypes than the one observed in single KO (Yoshida et al., 1997). Furthermore, in our Emx2 ChIP-Seq data, Emx1 motifs were also present in the top 10 identified motifs. The fact that *Pbx1* shares some common peaks with Emx2 (Golonzhka et al., 2015; Ypsilanti et al., 2021) and that we also found in our RIME data peptide of Pbx2, a homolog of Pbx1 known to cooperate with Pbx1 in the telencephalon (Golonzhka et al., 2015) and of Meis2, an evolutionarily conserved interactor of Pbx family members present in a similar gradient in the telencephalon as that of Pbx1 (Toresson et al., 2002b; Schulte & Geerts, 2019) also suggest it may be an Emx2 cooperative partner. However, the conditional knockout of *Pbx1* has been shown to reduce rostral areas, a phenotype opposite to that of *Emx2*, suggesting that they may antagonize each other’s function (Golonzhka et al., 2015).

**Figure 5:**
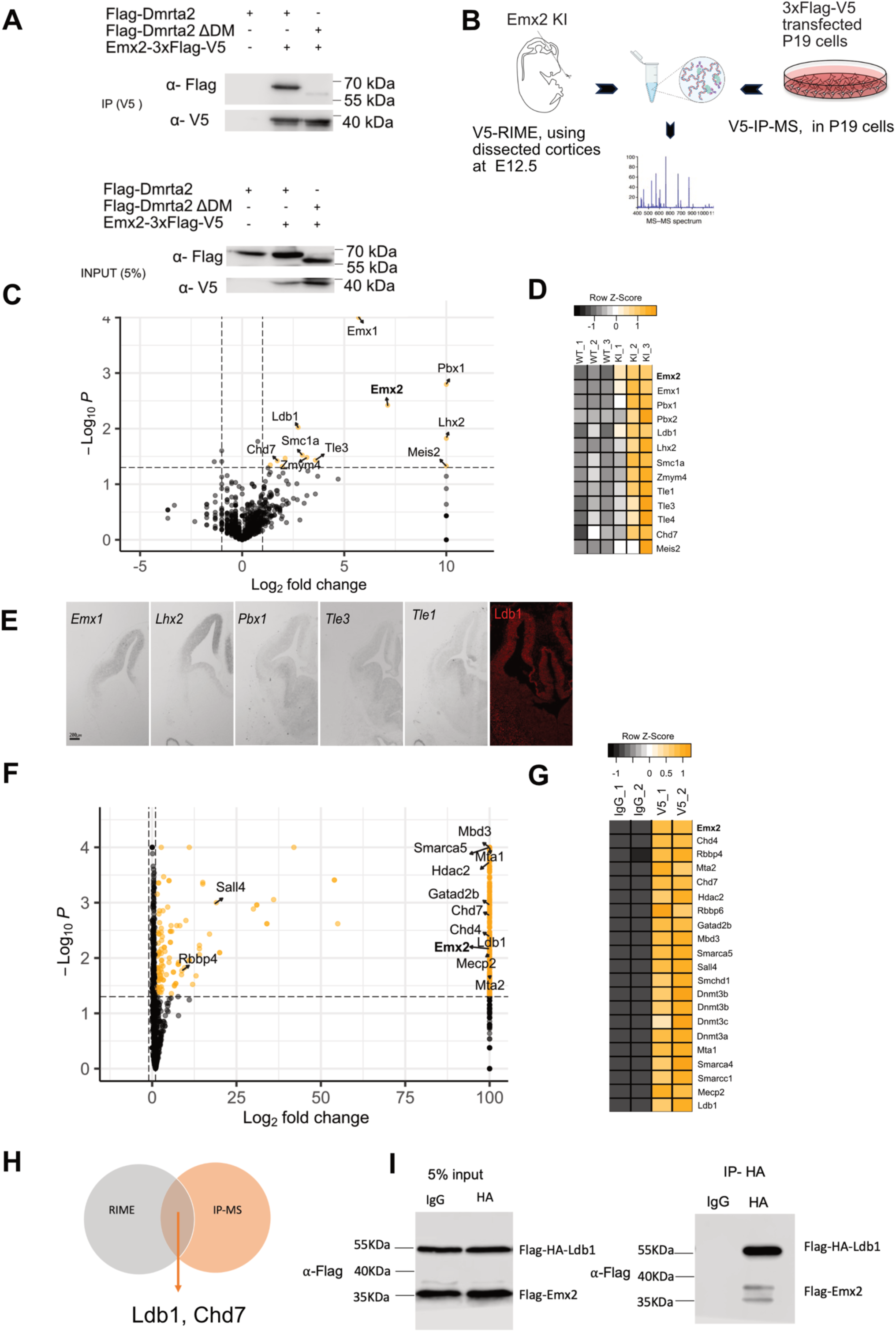
Identification of Emx2 interacting partners in embryonic dorsal telencephalic tissue. **A**. A co-immunoprecipitation assay with Flag-Dmrta2, Flag-Dmrta2ýDM and Emx2-3xFlag-V5 constructs overexpressed in HEK293T cells showing that Flag-Dmrta2 but not Flag-Dmrta2ýDM interact with Emx2 (n=3). **B**. Strategy for the identification of global interactive protein partners of Emx2. Rapid Immunoprecipitation of Endogenous proteins (RIME) was performed using V5 antibodies and E12.5 dissected cortices of Emx2 KI embryos. Immunoprecipitation followed by mass spectrometry (IP-MS) was performed using V5 antibodies and protein extracts from P19 cells transfected with an Emx2-3xFlag-V5 expression construct. **C**. A volcano plot showing the identified protein partners of Emx2 using RIME. All proteins that showed significant enrichment of total peptides are represented as orange-coloured points. Non-significant proteins are indicated as black points. Proteins that show only enrichment in the experimental (KI) sample compared to control (WT) samples (infinite enrichment) were given an arbitrary Log_2_(Fold Change) value of 10. No fold change cut-off was considered. Proteins of interest are indicated with name (n=3 for each condition). **D**. A heatmap showing the significantly enriched proteins identified by RIME and their normalised total spectral count across indicated samples (n=3 for each condition). **E**. Cross sections through the telencephalon of E12.5 embryos processed by ISH or IF showing the expression profile of the indicated identified Emx2 putative interactors. **F**. A volcano plot showing identified Emx2 interacting proteins using IP-MS. All proteins that showed significant enrichment of total peptides are represented as orange-coloured points. Non-significant proteins are indicated as black points. Proteins that show only enrichment in the experimental sample(V5) compared to control (IgG) samples (infinite enrichment) were given an arbitrary Log_2_(Fold Change) value of 100. Proteins of interest are indicated with name (n=2 for each condition). No fold change cut-off was considered. **G**. A heatmap showing the significantly enriched proteins identified by IP-MS and their normalised total spectral count across indicated samples (n=2 for each condition). **H**. A Venn diagram showing Ldb1 and Chd7 as common enriched proteins in RIME and IP-MS experiments. **I**. A co-immunoprecipitation assay performed with HEK293T cells overexpressing Flag-HA-Ldb1 and Flag-Emx2 constructs as indicated showing their interaction (n=3).

Besides TFs, peptides encoding cofactors were also enriched in Emx2 KI samples. Among them Tle cofactors that are mammalian homologs of *Drosophila* Groucho. Tle cofactors have been previously shown to interact with Emx2 *Drosophila* homolog ‘*Empty Spiracles* via its eh1 motif (Goldstein et al., 2005). They are coexpressed with Emx2, with *Tle1* showing expression in the hem and *Tle3* in the entire telencephalic ventricular zone. The LIM domain binding protein 1 (Ldb1) is another cofactor that we found enriched in Emx2 KI samples. Ldb1is ubiquitously expressed throughout the embryonic forebrain and is required for dorsal telencephalon development (Kinare et al., 2019).

In our IP-MS experiments on P19 cells transiently overexpressing a tagged version of Emx2, we also obtained a good enrichment of Emx2 peptides and of additional candidate interactive partners. Among them Ldb1 and many chromatin remodelling factors, including all components of the nucleosome remodelling and deacetylase (NurD) complex (**Fig. 5 F, G**). Combining RIME and IP-MS data sets allowed us to select two proteins as strong candidate Emx2 cofactors (**Fig. 5H**). The first one is the chromatin remodeler chromodomain helicase DNA-binding 7 (CHD7). CHD7 may be a relevant cofactor for Emx2 as it is expressed as early as E8.5 in the developing telencephalon (Jiang et al., 2012). The second and, compared to CHD7, the most enriched one is Ldb1, a well-known interactive partner of the Lim homeodomain family of transcription factors, including Lhx2. The fact that no Lim proteins were found in our IP-MS experiments suggests that Ldb1 faithfully interacts with Emx2 (Matthews & Visvader, 2003). To test this idea, we performed co-immunoprecipitation assays using tagged versions of Ldb1 and Emx2 overexpressed in HEK239T cells. Results obtained confirmed that they do interact (**Fig. 5 I**). Together, the data indicate that Dmrta2 is not a global cooperative factor of Emx2 and suggest that Emx2 interacts with the adaptor Ldb1 protein in telencephalic progenitors.

### Emx2 recruits Ldb1 to activate and repress its downstream targets

How Emx2 functions as a repressor is unknown. It may do so by forming a complex with Tle3, a known evolutionally conserved co-repressor, but Tle3 expression only starts at E12.5 while many of the defects of *Emx2* KO are already visible before that stage (Muzio et al., 2002b; Chytoudis-Peroudis et al., 2018). Alternatively, Emx2 may also directly recruit the NuRD complex component RBBP4 via a lysine and arginine-rich motif immediately preceding the homeodomain, as observed for other TFs including the Dlx1 homeobox TF (Price et al., 2022). However, using co-immunoprecipitation assays, we found that they do not interact (**Extended Data** Figure 6-1).

We then looked for the role of Ldb1 in the Emx2 function. Ldb1 is a known looping factor that binds to homeodomain and other types of TFs and helps them establish long-range enhancer-promoter contacts across diverse developmental pathways (Ma et al., 2008; Song et al., 2010; Zhang et al., 2015; Magli et al., 2019; Yasuoka & Taira, 2021). This is of particular importance in the developing nervous system in which many of the largest genes are expressed (McCoy & Fire, 2020; McCoy & Fire, 2024). In the developing cortex, Ldb1 has been shown to interact with the Lim homeodomain TF Lhx2. In agreement, the early loss of *Ldb1* largely phenocopies the defects observed in *Lhx2* mutants, with however some disparity in the expression of some hem-specific genes such as *Wnt2b*, suggesting Ldb1associates with other TFs (Mangale et al., 2008; Kinare et al., 2019). We hypothesised that if Ldb1 contributes to genes activated by Emx2, those genes would be expressed along a gradient similar to that of *Emx2*. Emx2 would bind both at the promoter and enhancer of these genes and they would be downregulated in *Emx2* mutants. To test this hypothesis, we chose two genes, *Nfib* that has been shown previously to be a direct target of Emx2 (Lim et al., 2021) and *Dach1*, that met these criteria. Previous PLAC-Seq experiments suggested that Emx2 bound enhancers in *Nfib* and *Dach1* do establish long-range contact with their promoters (Ypsilanti et al., 2021). *Dach1* particularly has been shown to have long-range enhancers located in “gene deserts” (Nobrega et al., 2003). We compared *Dach1* and *Nfib* expression in *Emx2^-/-^* and WT control embryos by ISH and RT-qPCR. Results in **Fig. 6A, C** show that their expression is indeed reduced in *Emx2* mutants. For both genes, we then performed ChIP-qPCR using Ldb1 antibodies and primers taken within the identified Emx2 bound putative enhancer and promoter regions, and in another region showing no Emx2 binding as a negative control. Results obtained shows that Ldb1 is recruited to both Emx2 bound enhancer and promoter regions of *Dach1*. For *Nfib*, a Ldb1 recruitment was observed on the promoter, but not the enhancer (**Fig. 6B, D**).

**Figure 6:**
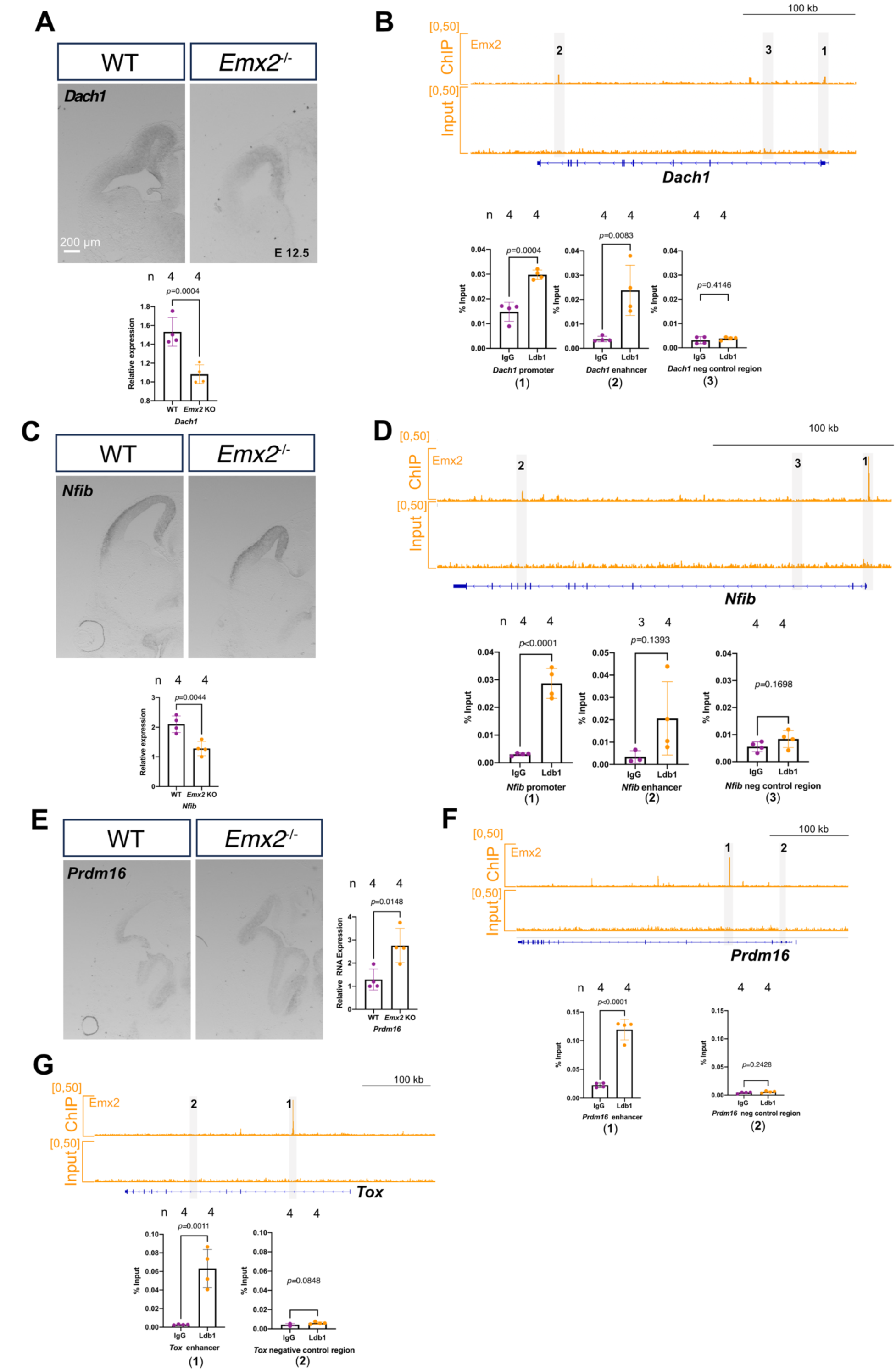
Emx2 utilizes Ldb1 for its direct target gene activation and repression. **A.** (Left panels) Coronal sections through the brain of an E12.5 *Emx2^-/-^* and WT embryos processed by ISH showing that *Dach1* and *Nfib* expression is reduced in the absence of *Emx2.* RT-qPCR on RNA extracted from dissected cortices quantifying this reduction is shown below. (Right panels) IGV snapshot showing Emx2 peaks at the promoter and in a putative enhancer region within the *Dach1* and *Nfib* locus and ChIP-qPCR data showing the level of enrichment of Ldb1 at the *Dach1* promoter (1), enhancer (2) and in an unbound region used as a negative control (3). **B**. (Left panel) Coronal sections through the brain of an E12.5 *Emx2^-/-^*and WT embryos processed by ISH showing that *Prdm16* expression is upregulated in the absence of *Emx2* and qRT-PCR recapitulating the same. (Right panels) IGV snapshot showing Emx2 peaks at the enhancer of *Prdm16* and Tox and ChIP-qPCR data showing selective enrichment of Ldb1 at *Prdm16* and Tox enhancers (1) and in unbound regions taken in those two loci used as negative controls (2). For all ISH n=3, For all qRT-PCR and ChIP-qPCR n=4. Error bars showing mean ± SD. p-value was calculated using an unpaired two-tailed T-test, (scale 200μm).

Ldb1 has been shown to repress downstream targets in a context-dependent manner, via the recruitment of the NurD complex through its interaction with MTA1,2 (Kiefer et al., 2011; Zhang et al., 2015). We hypothesised that this property of Ldb1 might contribute to the repression of Emx2 target genes. We tested this idea with two genes that are upregulated in *Emx2* KO embryos, *Prdm16* and *Tox*. We validated *Prdm16* upregulation in *Emx2* KO by ISH and RT-qPCR (**Fig. 6E**). We subsequently performed Ldb1 ChIP-qPCR on enhancers of *Prdm16* and *Tox* bound by Emx2. Our experiments show that they are also bound by Ldb1(**Fig. 6F, G**). Finally, to enquire about the association between NurD complex components and Ldb1, we immunoprecipitated Ldb1 in E12.5 cortical lysates. The results obtained show that Ldb1 interact with the NurD complex components MTA1 and HDAC2 (**Extended Data** Figure 6**-2**). These results suggest that Ldb1contributes to Emx2 activation and repression functions.

### *Dmrta2* is expressed in the developing choroid plexus, and its loss compromises choroid plexus functionality postnatally

The choroid plexus tissue is an epithelio-endothelial convolute responsible for producing the majority of the cerebrospinal fluid (CSF) (Solár et al., 2020). The dorsal telencephalic choroid plexus forms amid the choroid plaque and the cortical hem after the evagination of the dorsal midline (Kompaníková et al., 2022). In the hem region, whereas *Emx2* is expressed in the hem but is not detected in choroid plexus progenitors, *Dmrta2* is detected in the cortical hem and the developing choroid plexus (Tole et al., 2000; Saulnier et al., 2013). This is also evident in recent single-cell RNA-Seq experiments performed on telencephalic medial progenitors (**Extended Data** Figure 7-1 **A**) (Moreau et al., 2023). Early electroporation of *Emx2* in the chicken choroid plexus epithelium suppresses choroid plexus development and promotes neuroepithelial cell fate (von Frowein et al., 2006). To uncover the role played by *Dmrta2* in this tissue, we conditionally ablated *Dmrta2* using the *Msx1:Cre-ERT2* line that selectively ablates *Dmrta2* in the choroid plexus progenitors. First, to confirm that this line can drive Cre-mediated recombination in the developing choroid plexus tissue, we crossed it with the *Rosa-Stop-YFP* line. Tamoxifen was injected intraperitoneally into pregnant *Msx1Cre-ERT2; Rosa-Stop-YFP* dam at E9.5. Embryos were harvested at E12.5 and analyzed by immunostaining for YFP expression. In those embryos, YFP expression similar to *Msx1* expression was observed in the choroid plexus and in the meninges at E12.5, thus validating the line for the study of *Dmrta2* function in choroid plexus (**Extended Data** Figure 7-1 **B**). We then assessed the expression of the choroid plexus specification marker *Transthyretin* (*Ttr*) in both tamoxifen-injected *Dmrta2* icKO and WT embryos at E12.5. No change in *Ttr* expression was observed in these *Msx1Cre-ERT2;Dmrta2^fl/fl^*compared to control embryos (**Extended Data Figgure 7-1C)**. As embryonic tamoxifen administration can adversely affect *Dmrta2* expression in WT (Lee et al., 2020), we did not use further these *Msx1Cre-ERT2;Dmrta2^fl/fl^*mice and turned to the *Lmx1a-*Cre line, which expresses Cre both in the choroid plexus and the cortical hem starting from E10.5 (Parichha et al., 2022). In *Lmx1aCre;Dmrta2^fl/fl^*, henceforth *Dmrta2* cKO, *Dmrta2* expression was lost in the cortical hem and choroid plexus, as evidenced by ISH analysis of *Dmrta2* expression using an antisense probe against exon 2. An upregulation of the *Dmrta2* was observed using a probe against exon 3 in the choroid plexus of *Dmrta2* cKO embryos, as previously observed in the cortex, likely due to *Dmrta2’s* ability to negatively regulate its own expression in a feedback loop (De Clercq et al., 2018) (**Fig. 7A**). In these *Dmrta2* cKO embryos, we examined *Wnt* expression in the cortical hem, given its importance for choroid plexus development (Lee et al., 2000; Parichha et al., 2022). We observed that the expression of *Wnt3a,* the earliest Wnt gene expressed selectively in this region appears similar in *Dmrta2* cKO embryos than in controls. In *Dmrta2* cKO embryos, *Lmx1a* expression in the cortical hem and in the choroid plexus appears also normal. There was also no change in the expression of *Msx1* or *Ttr* that both mark the developing choroid plexus epithelium. Thus, *Dmrta2* does not appear to be required for the specification of the telencephalic choroid plexus. However, we found that *Dmrta2* cKO animals developed hydrocephalus and severe ataxia postnatally with growth retardation and enlarged ventricles characteristic of hydrocephaly (Dietrich et al., 2009; Yang et al., 2019) (**Fig. 7B-D**). Hydrocephalus is often linked with defective choroid plexus formation or functionality (Makiyama & Mizusawa,1997; Vong et al., 2021; Yang et al., 2019). We therefore subjected these mice for [^18^F]-fluorodeoxyglucose ([^18^F]-FDG) imaging by positron emission tomography (µPET) combined with Computed Tomography(µCT) to substantiate the hydrocephalus phenotype. Our PET combined CT analysis confirmed that cKO possess larger fluid-filled ventricles with metabolically inert or absent areas compared to WT mice brains (**Fig. 7E**). Mean Standardized Uptake Value (SUVmean) associated with the Volumes Of Interest (VOIs) was drawn to encompass the brain’s lower radioactive glucose uptake in those areas in cKO compared to WT mice brains. We further analyzed various markers of junctions and lamina of the blood CSF barrier and the choroid plexus epithelia to understand the reasons for such a phenotype. The markers examined, including laminin, ZO-1 and β-Catenin showed no difference in expression or localisation in the telencephalic choroid plexus epithelia between cKO and controls (**Extended Data** Figure 7-2 **A-C)**.

**Figure 7:**
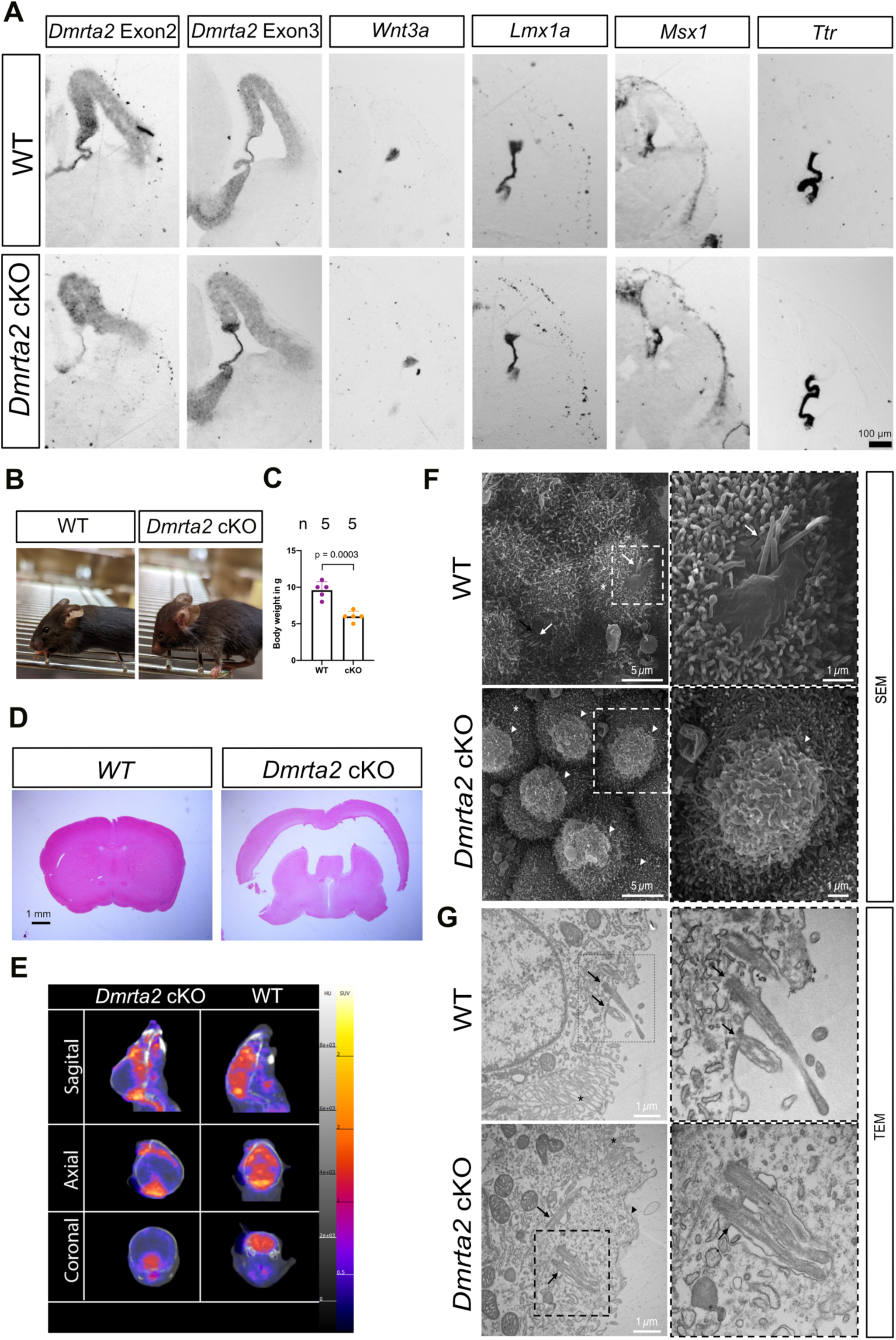
Conditional ablation of *Dmrta2* in the cortical hem and choroid plexus does not impair choroid plexus specification but leads to hydrocephaly. **A.** Cross-sections through the telencephalon of E12.5 *Dmrta2* cKO and WT controls processed by ISH for the indicated probes showing that the loss of *Dmrta2* does not alter the expression of early choroid plexus markers (n=3) (scale bar 100μm) **B**. Photograph of a WT mouse at P21 and of a littermate *Dmrta2* cKO showing a dome-shaped head characteristic of hydrocephaly. **C**. Graph showing the reduction in weight of *Dmrta2* cKO compared to WT mice at P21. (WT= 4, cKO= 5). Error bars showing mean ± SD. p-value was calculated using an unpaired two-tailed T-test, **D**. Hematoxylin-eosin stained cross sections of the brain of *Dmrta2* cKO and WT control mice showing that *Dmrta2* cKO mice have enlarged ventricles characteristic of hydrocephaly (scale bar 1mm). **E**. Representative sagittal, coronal and axial images showing reduced emission intensity in the *Dmrta2* cKO head compared to wild-type animals of P21-P30 age (WT=4, cKO=3). The colour scale bar shows the scale of SUV value; the greyscale bar shows the scale of HU (Hounsfield unit). **F, G**. SEM(F) and TEM(G) images showing that the morphology of the telencephalic choroid plexus apical cell membrane is altered in *Dmrta2* cKO mice. Note in WT mice the presence of epithelial cells covered with microvilli (asterisks) with patches of short cilia (arrows) emerging from flat regions of the plasma membrane. In the ChP of Dmrta2 cKO mice, protrusions were regularly observed on the top of the epithelial cells (arrowheads). Those masses directly emerge from the cytoplasm and are located in close vicinity to the cilia.

To examine the ultrastructure of the telencephalic choroid plexus epithelium, we dissected the telencephalic choroid plexus from both control and Dmrta2 cKO mice and subjected the tissue to Scanning Electron Microscopy (SEM) and Transmission Electron Microscopy (TEM) imaging. SEM images of WT choroid plexus revealed cells mostly covered with microvilli, alongside patches of short cilia at the cell apex. These ciliary structures were often associated with flat regions of the plasma membrane, which sometimes exhibited small protrusions of varying shapes (**Fig. 7F**). TEM analysis confirmed the presence of short, non-motile (9+0 axoneme) cilia in specialized membrane regions devoid of microvilli. In contrast, SEM images of cKO choroid plexus showed numerous cells covered by masses with shapes and sizes inconsistent with typical immune cells found in direct tissue contact (**Fig. 7G**). TEM sections verified that these masses showing irregular morphologies originate from epithelial cells and are filled with impoverished cytoplasm, lacking mitochondria and containing only minimal endoplasmic reticulum (**Extended Data** Figure 7-3A). However, ribosome-like and vesicle-like structures, as well as multilamellar bodies, were visible. Notably, cilia were regularly observed in close vicinity to the masses, suggesting their origin from specialized flat membrane regions associated with ciliary structures in the choroid plexus. Overall, apart from these masses, the tissue appeared healthy in SEM and TEM observations, with highly active epithelial cells displaying densely stained mitochondria and exhibiting normal cell-to-cell junctions (**Extended Data** Figure 7-3B). These observations hint at the possibility that the specialized plasma membrane associated with cilia may lose its rigidity, leading to the formation of protrusions on the cell apex. These modifications could potentially impact or be a consequence of alterations in ciliary functions. Although it cannot be ruled out, no evident defect of the ciliary structure could, however, be observed in this study. In SEM, we also observed otherwise normal ependymal cilia (**Extended Data** Figure 7-3C).

A recent study has shown that increased intracellular Ca^2+^ signaling can lead to apocrine secretion-related protrusion in ChP (Courtney et al., 2024). We, therefore, performed c-Fos staining to assess Ca^2+^ signaling (Sheng et al., 1900) in ChP but observed no significant change in c-Fos staining between *Dmrta2* cKO and controls (**Extended Data** Figure 7-4). In conclusion, although *Dmrta2* is dispensable for the specification of the choroid plexus, it appears to be required for its functionality in adult mice.

## DISCUSSION

*Dmrta2* and *Emx2* are two genes expressed similarly in the developing telencephalon since its formation. Their individual loss has been shown to impart a wide range of similar phenotypic changes across various developmental stages of the cortex (Yoshida et al., 1997; Tole et al., 2000; Mallamaci et al., 2000; López-Bendito et al., 2002; Mallamaci, 2011; Konno et al., 2012; Saulnier et al., 2013; Falcone et al., 2015; Muralidharan et al., 2017; Ratié et al., 2020) and their compounded deletion to exacerbate the phenotypes (Desmaris et al., 2018) suggesting possible cooperation during cortex development. The mechanism behind this cooperation remains unknown.

In this study, we show that the cerebral hemispheres are absent in *Emx2^-/-^;Dmrta2*^-/-^ E18.5 embryos, which further highlights the importance of their cooperation for corticogenesis. Pallial markers analyzed at E12.5 in double knockout indicate near complete loss of all markers, similar to the loss of *Ngn2* we observed previously (Desmaris et al., 2018). On the other hand, we observed an up-regulation of subpallial markers, suggesting a transformation of pallium into subpallium laterally. Interestingly, similar observations were made in the *Emx2;Pax6* double knockout (Muzio et al., 2002a; Kimura et al., 2005). In *Emx2;Pax6* double knockout, though the pallium is specified, it is subsequently transformed into subpallium, whereas the medial pallium is converted to a roof fate. A similar transformation might have occurred in *Dmrta2;Emx2* double knockout embryos. Though we have not analyzed roof markers in *Dmrta2;Emx2* double knockout embryos, it is likely the case, given the absence of all other medial markers. The upregulation of subpallial genes in *Dmrta2;Emx2* double knockout could be the result of the combined role of *Dmrta2* and *Emx2* in maintaining the pallium subpallium boundary (PSB) (Desmaris et al., 2018). The changes in medial pallium transformation can result from coordination between telencephalic and diencephalic transcription factors such as Otx1,2 in establishing the caudal forebrain primordium (Kimura et al., 2005). *Emx2* has been shown to play a role in this regard, but *Dmrta2*’s role remains to be explored (Suda et al., 2001).

Our bulk RNA sequencing of Emx2 knockout telencephalon and subsequent comparison with *Dmrta2* knockout bulk RNA sequencing data reinstates similar roles played by both TFs during early cortical development, as suggested by earlier studies (Saulnier et al., 2013). The similarity of phenotypes and deregulated genes upon the loss of *Dmrta2* or *Emx2* suggests they likely have many common direct targets. Unexpectedly, our comparative ChIP-data analysis of Emx2 and Dmrta2 yielded only a few common direct downstream targets. Most genes bound by Emx2 and Dmrta2 were not expressed in the medial pallium, where their expression is high. In agreement with our analysis, a recent study that combined the ChIP-Seq data set of Emx2, Pbx1, Couptf1, Pax6 and Lhx2 in the dorsal telencephalon has found that they all bind to less number of genes expressed in the medial pallium than in the lateral pallium (Ypsilanti et al., 2021). Even though Emx2 and Dmrta2 had only a few common direct targets, among them were essential determinants of alternate cortical fate. We subsequently showed that the expression pattern of these cortical fate determinants is affected in *Emx2* and *Dmrta2* single and double knockouts. Gene expression changes were subtle in Emx2 single knockouts compared to *Dmrta2* single knockouts. This aligns with the fact that Dmrta2 can regulate *Emx2* and impart a more severe phenotype upon its loss (Tole et al., 2000; Saulnier et al., 2013). Though they lack many common direct targets, regulating key cortical fate determinants might be sufficient to impart similar gene expression program changes and generate a similar phenotypic output upon their loss.

Our Co-IP assays indicated that Emx2 and Dmrta2 can interact and that this interaction is compromised upon loss of the DM domain of Dmrta2. This suggest that the DM domain of DMRT proteins that binds to DNA and is required for dimerization (Murphy et al., 2007; Murphy et al., 2015) could be also involved in the association with other proteins. The fact that Emx2 and Dmrta2 only bind to a few common enhancers *in vivo* could arise because of a lack of adjacently placed TF motifs to stabilise their interaction on DNA. To identify Emx2 partners in a global manner, we resort to RIME using dissected cortices of E12.5 mouse embryos and IP-MS after transient transfection of Emx2 in P19 cells. Our results suggest that Emx2 cooperates with known cortical TF such as Emx1, Lhx2 and Pbx1 but not Dmrta2. No Emx2 enrichment was also observed in complementary Dmrta2 RIME experiments (X. Shen, unpublished observations), further suggesting the two cofactors cooperate on a limited set of targets. Emx1 is a cognate protein of Emx2, and genetic evidence suggests they cooperate during cortical development (Shinozaki et al., 2002; Bishop et al., 2002; Muzio et al., 2003; Shinozaki et al., 2004). Our results indicate they cooperate at the molecular level as well. *Lhx2*, on the other hand, is expressed in a similar gradient as *Emx2* but is excluded from the hem. A recent study has shown that Emx2 and Lhx2 share the most enhancers in the dorsal telencephalon (Ypsilanti et al., 2021), validating our RIME approach for the identification of novel Emx2 candidate interacting factors. Pbx1 is such a potential Emx2 interacting TF, given their demonstrated cooperation during scapula development (Capellini et al., 2010). It is however expressed in an opposite gradient of Emx2, and genetic evidence suggests that their interaction is antagonistic (Golonzhka et al., 2015). We also identified candidate Emx2 cofactors such as Tle3 and Ldb1 in our RIME experiments. However, *Tle3* expression in the dorsal telencephalon starts at E12.5, whereas Emx2 loss of functions phenotypes are already visible before E12.5, suggesting this interaction might be important for the later neurogenic phase (Chytoudis-Peroudis et al., 2018). A distinct set of interacting factors has been recently identified in similar Dmrta2 RIME experiments performed in parallel to the Emx2 RIME experiments described in this study (X. Shen, unpublished observations), suggesting that the mechanisms used by Emx2 and Dmrta2 to regulate their targets are largely distinct.

We used IP-MS data from P19 cells to narrow down the potential physiologically relevant partners obtained through RIME. Combining our RIME and IP-MS data sets, we identified Ldb1 and Chd7 as interactive partners of Emx2 in both conditions, though peptide counts were low for Chd7. Ldb1 is a well-known interactive partner of Lim homeodomain transcription factors, but at least in our IP-MS data, we did not find any such proteins (Matthews & Visvader, 2003). Hence, we further explored the role of Ldb1 as an interactive partner of Emx2. It is, however, important to note that loss of function of Chd7 leads to a similar medial defect to that of *Emx1;Emx2* knockout (Jiang et al., 2012). We validated this interaction further by Co-IP and demonstrated that Ldb1 might contribute to activating and repressing roles of Emx2 in some of its downstream targets. We cannot, though, exclude that Lhx2 might as well recruit Ldb1 to these enhancers as most of them are bound by Lhx2 (Ypsilanti et al., 2021).

In contrast to *Emx2* that has been implicated in the repression of ChP (Von Frowein et al., 2006), *Dmrta2* is expressed in the medial pallium in choroid plexus progenitors. Later, Dmrta2, like Emx2, remains enriched in differentiated telencephalic compared to hindbrain choroid plexus tissue (Lun et al., 2015). We thus also interrogate in this study whether *Dmrta2* plays any role in its development. To this end, we utilised the *Msx1:Cre^ERT2^* line that selectively ablates *Dmrta2* in the choroid plexus progenitors or the *Lmx1a*-Cre line that ablates *Dmrta2* in the hem and choroid plexus. We did not observe any specification defects in either case. *Lmx1a*-Cre conditional knockout animals however postnatally developed hydrocephaly suggesting a compromised choroid plexus function. We speculate that this phenotype reflects a direct consequence of the loss of *Dmrta2* in ChP rather than an indirect consequence of poorly developed WNT-rich cortical hem known to be required for ChP development (Lee et al., 200; Parichha et al., 2022) as we did not observe alteration of the expression of *Wnt3a*. In *Wnt3a*-Cre; *Dmrta2^fl/fl^* embryos in which *Dmrta2* is deleted specifically in the hem from E10, Wnt signaling has been shown to be also unaffected (De Clercq et al., 2018). Analysis of the ultra-structure of the telencephalic choroid plexus epithelium using SEM and TEM revealed that this phenotype is not a consequence of compromised epithelial junctions or defects in ependymal cilia (Narita & Takeda, 2015). We however observed that the apical surface of the choroid plexus epithelium of *Dmrta2* cKO mice possesses protrusions or masses. These protrusions were often observed near ChP cilia, suggesting their role in the compromised cytoarchitecture. We also observed stenosis of the aqueduct in the cKO mice (**Extended Data** Figure 7-5), which could potentially contribute to the observed hydrocephalic phenotype (Wagner et al., 2003; Agerskov et al., 2020; Roales-Buján et al., 2021; Rados et al., 2021) observed in *Lmx-Cre;Dmrt5^fl/fl^* mice. Indeed, *Lmx1a* is co-expressed with *Dmrta2* in the ventral midbrain. The Cre recombinase is thus likely also active in this brain region in which *Dmrta2* has been suggested to plays role in its patterning, as revealed by an *in vitro* study using ES cells differentiated into ventral midbrain neurons (Gennet et al., 2011). We consider this scenario however unlikely given the fact that, in contrast to what has been observed *in vitro*, *Dmrta2* inactivation *in vivo* does not affect ventral mesencephalic neural fate specification (Sirakov et al., personal communication).In conclusion, our study provides novel insights into the way Emx2 cooperates with Dmrta2 to control telencephalon patterning. It also provides the first evidence suggesting that Dmrta2 plays a role in the ChP function, which remains to be further explored.

## Author contributions

J.A. and E.J.B. designed the experiments. J.A., X. S., T.S., L.C., and G.D. performed the experiments. J.A., X. S., T.S., M.D., A.A., L.C., G.D, M.S., G.S.R., P.R. analysed the data. Y.S. and S.K. generated reagents. J.A. and E.J.B. wrote the manuscript.

## Acknowledgements

This work was supported by the FNRS (CDR J001121F to EJB). JA was a FNRS-FRIA fellow and thereafter obtained a “Fonds David and Alice Van Buuren” fellowship. XS was a China Scholarship Council fellow. TS was a ULB-UMons fellow. We thank Dr. Antonello Mallamaci for the kind gift of *Emx2* knockout mice, Dr. Kathleen J. Millen for the kind gift of *Lmx1a^Cre^* mice and Dr. Michel Cayouette for the kind gift of *Rosa-Stop-EYFP*; *Msx1:Cre^ERT2^* mice. We thank Dr Nicoletta Kessaris for the kind gift of the HA-Flag-Ldb1 plasmid construct. We thank Prof. David Pérez-Morga, Daniel Monteyne and Valérie Suain for the technical help with hydrocephalus SEM and TEM work. We thank Dr. Maxime Bellefroid, Dr. Sylvain Fauquenoy, and Louis Delhaye for their technical assistance. We want to thank Alba Sabaté San José, Dr. Maya Dannawi, and other members of the laboratory of Developmental Genetics for their helpful discussion.

