## Extended Data Figures and Table for "Novel insights into Emx2 and Dmrta2 cooperation during cortex development and evidence for Dmrta2 function in choroid plexus"

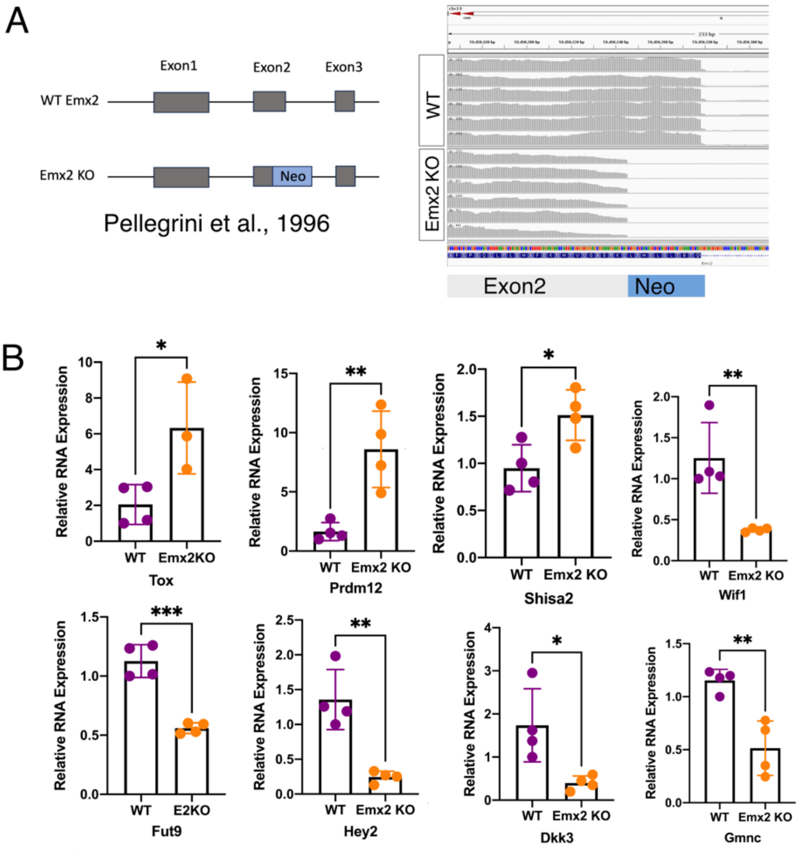


**Extended Data Figure 1-1. Validation of the Emx2 KO line and RT-qPCR confirming the deregulation of selected genes identified by bulk RNA-seq of the transcriptome of dorsal telencephalic tissue dissected from E12.5 *Emx2* knockout embryos. A** . WT vs *Emx2* knockout alleles as per Pellegrini et al., 1996**.** IGV view of the bulk RNA-seq data at the Emx2 locus obtained examining 5 KO and 5WT samples. Note that, as expected, no transcript corresponding to the end of exon 2, replaced by *neo*, has been sequenced in the KO samples. **B.** RT-qPCR showing relative RNA expression of indicated genes in *Emx2* KO versus WT. For each case n=4 for WT and KO, except for *Tox*, where n=3 for KO. An unpaired two-tailed T-test was used to infer p-value. n represents biologically independent replicates. ***(P<0.001), **(P<0.01), *(P<0.05).

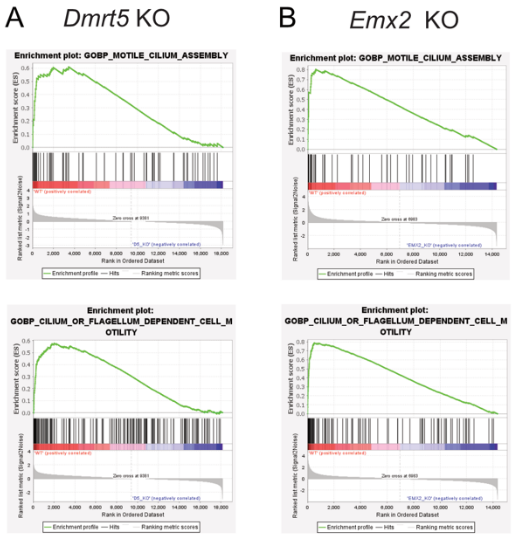


**Extended Data Figure 2-1. Gene sets related to motile cilium assembly and cilium and flagellum-dependent cell mobility are downregulated in *Dmrta2* and *Emx2* knockouts**. Enrichment plots of two common gene sets enriched in GSEA analysis of *Dmrta2* KO (A) and *Emx2* KO(B) bulk RNA-Seq data, showing the profile of the running ES Score and positions of gene set members on the rank-ordered list.

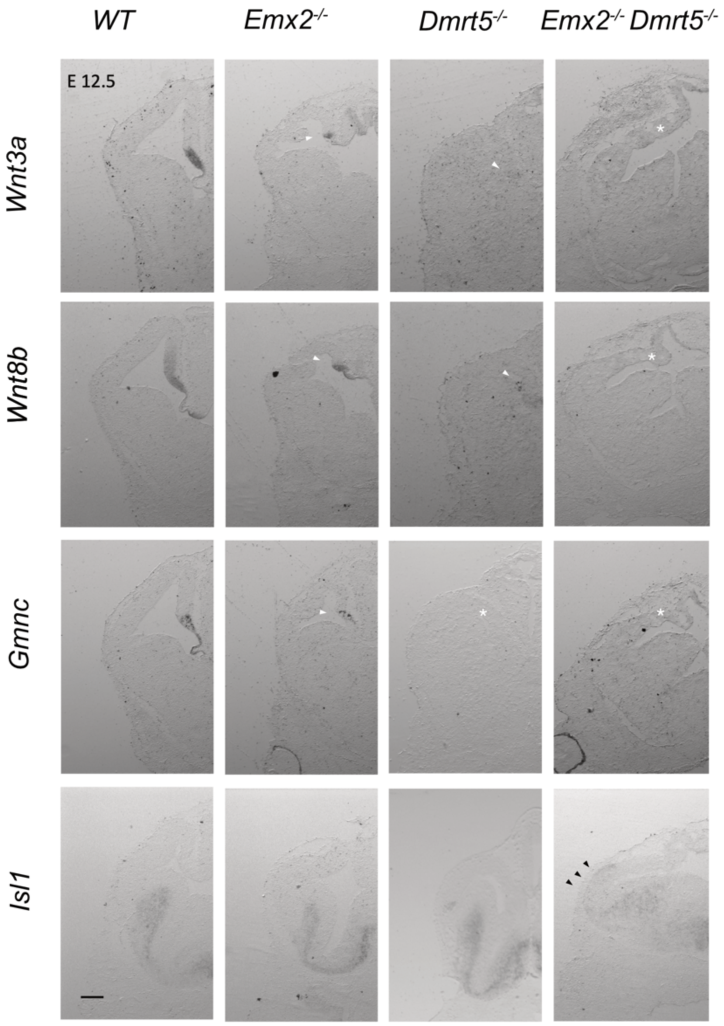


**Extended Data Figure 2-2. Dorsal telencephalic markers are downregulated and the Islet1 vLGE marker expands in the dorsal telencephalic neuroepithelium of *Emx2;Dmrta2* double KO embryos**. Cross-sections of the telencephalon of E12.5 embryos of the indicated genotypes (n=3) processed by ISH to reveal the expression of the indicated genes are shown. White arrowheads show decreased gene expression, white asterisks indicate loss of gene expression, and black arrowheads indicate increased gene expression. Scale bar 100µm.

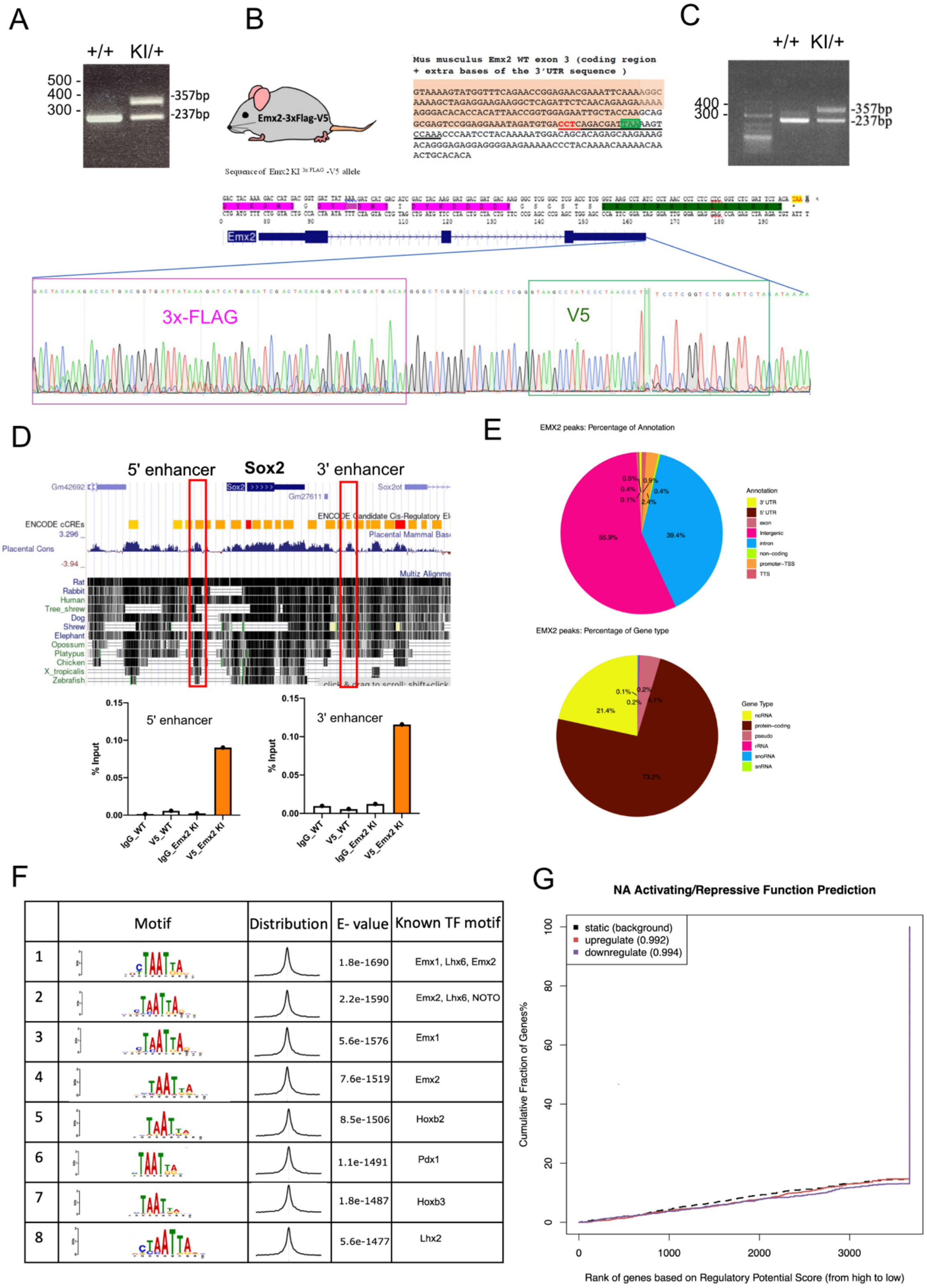

A
+/+
KI/+
500 -
400 -
300 -
-357bp
-237bp
B
Emx2-3xFlag-V5
Sequence of Emx2 KI 3x FLAG -V5 allele
Mus musculus Emx2 WT exon 3 (coding region + extra bases of the 3'UTR sequence )
GTAAAAGTATGGTTTCAGAACCGGAGAACGAAATTCAAAAGGC
AAAAGCTAGAGGAAGAAGGCTCAGATTCTCAACAGAAGAAAAA
AGGGACACACCACATTAACCGGTGGAGAATTGCTACCAAGCAG
GCGAGTCCGGAGGAAATAGATGTGACCTCAGACGATTAAAAGT
CCAAACCCAATCCTACAAAAATGGACAGCACAGAGCAAGAAAG
ACAGGGAGAGGAGGGGGAAGAAAAACCCTACAAAACAAAAACAA
ACTGCACACA
Emx2
3x-FLAG
V5
C
+/+
KI/+
400 -
300 -
-357bp
-237bp
D
5' enhancer
Sox2
3' enhancer
ENCODE cCREs
Placental Cons
Rat
Rabbit
Human
Tree_shrew
Dog
Shrew
Elephant
Opossum
Platypus
Chicken
X_tropicalis
Zebrafish
% Input
IgG_WT
V5_WT
IgG_Emx2 KI
V5_Emx2 KI
E
EMX2 peaks: Percentage of Annotation
Annotation
3' UTR
5' UTR
exon
Intergenic
intron
non-coding
promoter-TSS
TTS
55.9%
39.4%
EMX2 peaks: Percentage of Gene type
Gene Type
ncRNA
protein-coding
pseudo
rRNA
snoRNA
snRNA
21.4%
73.2%
F
Motif
Distribution
E- value
Known TF motif
1 1.8e-1690 Emx1, Lhx6, Emx2
2 2.2e-1590 Emx2, Lhx6, NOTO
3 5.6e-1576 Emx1
4 7.6e-1519 Emx2
5 8.5e-1506 Hoxb2
6 1.1e-1491 Pdx1
7 1.8e-1487 Hoxb3
8 5.6e-1477 Lhx2
G
NA Activating/Repressive Function Prediction
static (background)
upregulate (0.992)
downregulate (0.994)
Cumulative Fraction of Genes%
Rank of genes based on Regulatory Potential Score (from high to low)

**Extended Data Figure. 3-1. Generation and validation of a mouse knockin line expressing a C-terminal 3xFlag-V5 tagged version of the Emx2 protein (Emx2 KI) and some of the data obtained from the analysis of the V5 ChIP -seq experiments performed on Emx2 KI and WT as controls. A, B.** Genotyping followed by sequencing confirming the insertion of the 3xFlag-V5 tag at the c terminal end of the *Emx2* ORF (exon 3). In B, in the sequence of Emx2 exon3, the coding part is highlighted in pink, the PAM site for Cas9 is indicated in red, and the stop codon in green. **C**. RT-PCR on cDNA prepared from dissected cortices from a E12.5 heterozygote knockin and a WT control embryo using primers flanking the in sertion site. As expected, a 375 bp band corresponding to *Emx2* 3xFlag-V5 tag allele is only detected in the heterogygote KI sample. **D**. Representative ChIP-qPCR results obtained using V5 or IgG antibodies on chromatin prepared from cortices dissected from E12.5 Emx2 KI embryos or WT as controls and primers taken in conserved regions located 5' and 3' of the Sox2 genes (red boxes) acting as telencephalic enhancers bound and negatively regulated by Emx2 (Mariani et al., 2012). Note the enrichment of the tagged Emx2 protein on both enhancers, not detectable using IgG antibodies, or using V5 on WT controls. **E**. Emx2 peak annotation and associated gene type. **F**. Top 8 known TF motifs identified by MEME as enriched in Emx2 peaks. **G**. BETA analysis for Dmrta2 showing no significant association between peaks and deregulated genes.

| Protein | WT_1 | WT_2 | WT_3 | KI_1 | KI_2 | KI_3 |
|---|---|---|---|---|---|---|
| Emx2 | 0 | 2 | 1 | 95 | 123 | 130 |
| Emx1 | 0 | 0 | 1 | 10 | 21 | 17 |
| Pbx1 | 0 | 0 | 0 | 7 | 20 | 20 |
| Pbx2 | 0 | 0 | 0 | 0 | 6 | 10 |
| Ldb1 | 0 | 5 | 0 | 7 | 11 | 11 |
| Lhx2 | 0 | 0 | 0 | 4 | 14 | 20 |
| Smc1a | 0 | 16 | 3 | 13 | 68 | 66 |
| Zmym4 | 0 | 5 | 0 | 4 | 19 | 22 |
| Tle1 | 0 | 3 | 0 | 5 | 16 | 22 |
| Tle3 | 0 | 7 | 2 | 10 | 19 | 37 |
| Tle4 | 0 | 6 | 2 | 6 | 15 | 28 |
| Chd7 | 0 | 8 | 4 | 5 | 16 | 18 |
| Meis2 | 0 | 0 | 0 | 3 | 3 | 11 |

**Extended Data Table 5-1.** Table showing the total number of spectral counts corresponding to candidate interactive protein partners obtained in Emx2 RIME experimental replicates.

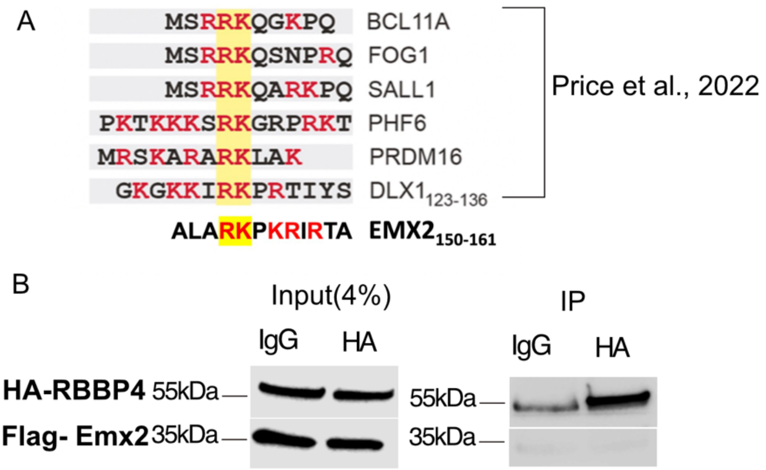


**Extended Data Figure 6-1. Emx2 does not interact with RBBP4 A.** Emx2 has a similar lysine and arginine-rich region as observed in other RBBP4-interacting proteins. B. A Co-IP assay using the indicated tagged proteins overexpressed in HEK293T cells showing that RBBP4 and Emx2 do not interact.

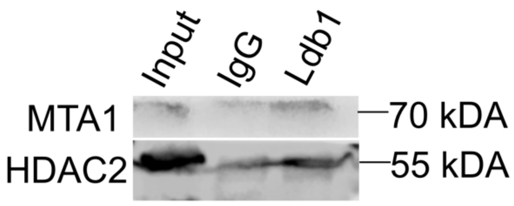


**Extended Data Figure 6-2. Ldb1 interacts with MTA1 and HDAC2. A Co-IP assay** using an Ldb1 antibody on E12.5 cortical lysate showing the interaction of Ldb1 with endogenous MTA1 and HDAC2 proteins.

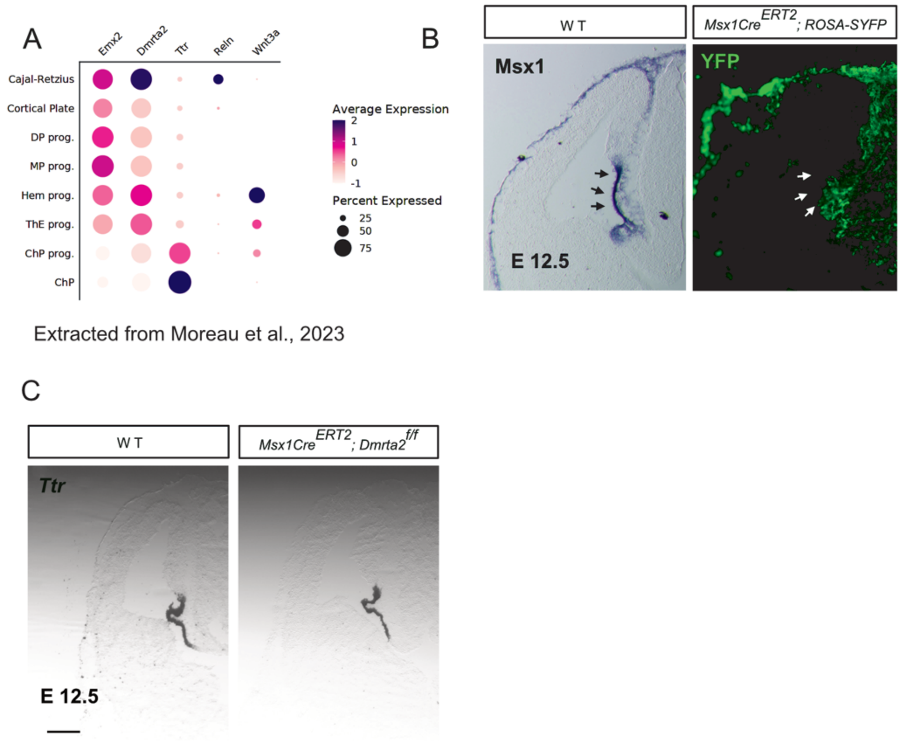


**Extended Data Figure 7-1. Dmrta2 expression is detected in chroïd plexus progenitors and its conditional abalation using the *Msx1Cre$^{ERT2}$* line does not affect the expression of the *Ttr* early choroid plexus marker. A**. Single-cell RNA-seq data from Moreau et al., 2023 showing the expression of *Dmrta2* and *Emx2,* together with *Ttr, Reelin* and *Wnt3a* as markers of progenitors of the choroid plexus, Cajal-Retzius cells and of the hem, respectively, in medial telencephalic progenitors (DP-Dorsal progenitors, MP- Medial progenitors, ThE- Thalamic Eminence, ChP- Choroid Plexus). ***B***. Cross-sections through the telencephalon of E12,5 *Msx1Cre$^{ERT2}$*;*ROSA-YFP* embryos injected with tamoxifen at E9,5 and processed by immunostained for YFP and by ISH for *Msx1* expression*,* showing that YFP is expressed similarly to Msx1. **C.** Cross-sections through the telencephalon of E12.5 *Msx1Cre$^{ERT2}$; Dmrta2$^{fl/fl}$* and WT embryos injected with tamoxifen at E9.5 processed by ISH for *Ttr* expression, showing it is unaltered in the absence of *Dmrta2*. Scale 100µm.

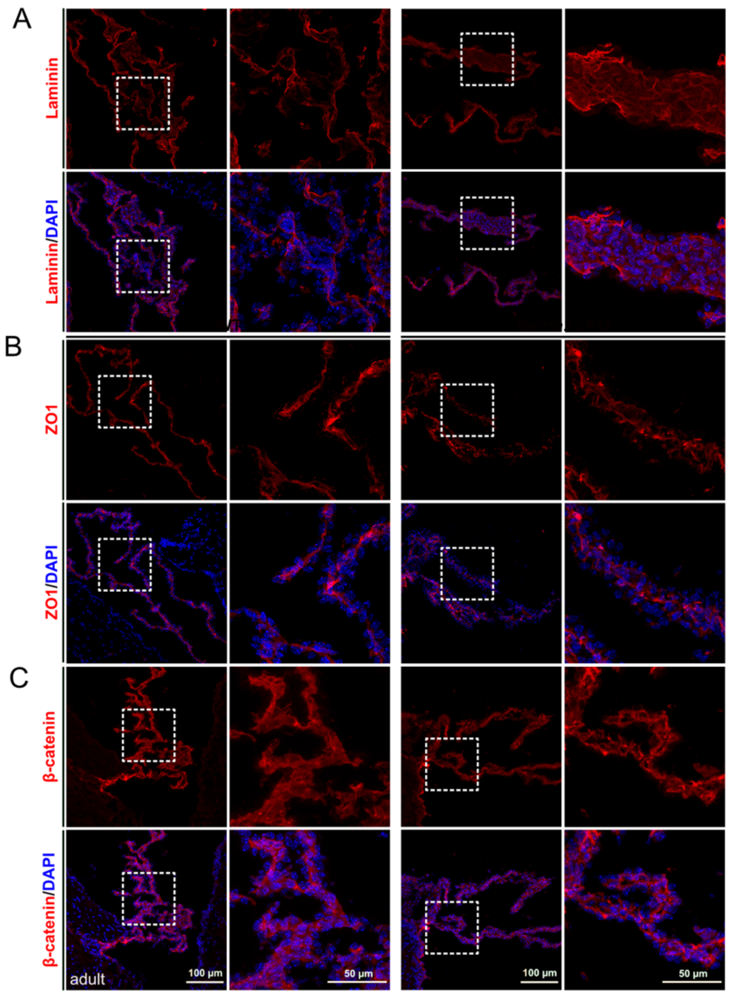


**Extended Data Figure 7-2. Cell junctions are intact in adult *Dmrta2* cKO epithelium. A, B.** IF staining of ZO-1, β-catenin showing intact adherens junctions in the adult *Dmrta2* cKO choroid plexus epithelium. **C.** IF staining of laminin showing intact lamina in the adult *Dmrta2* cKO choroid plexus epithelium.

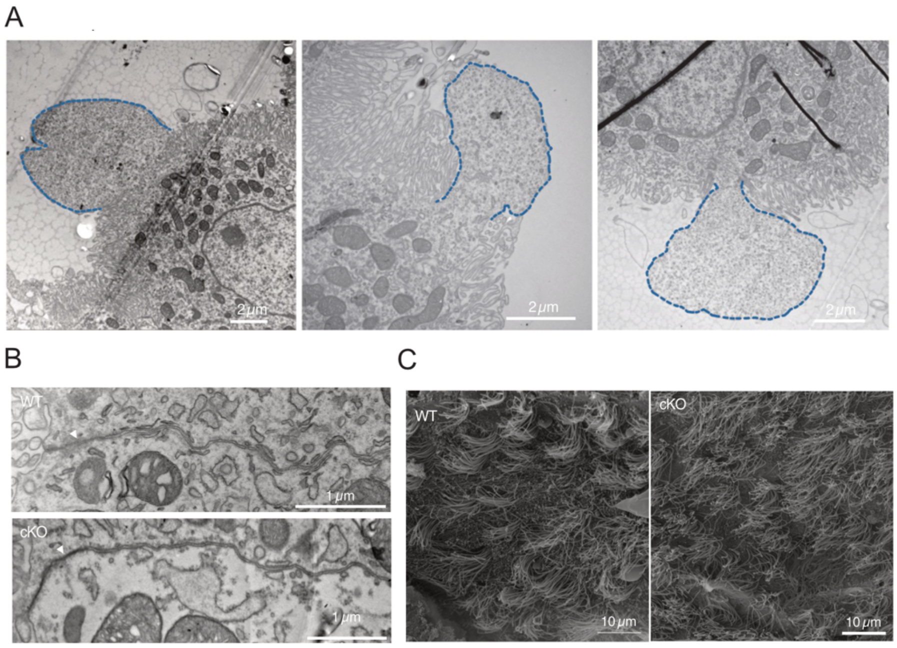


**Extended Data Figure 7-3. TEM and SEM observations of the choroid plexus epithelium of *Dmrta2* cKO mice. A.** TEM sections showing that ChP protrusions (dashed blue line) in *Dmrta2* cKO mice have irregular morphologies. They vary from semi-circular to irregular based on the plane of the section. **B**. TEM images showing normal tight junctions (arrowheads) in the ChP epithelia of cKO, as observed in WT mice. **C**. SEM images showing that ependymal cilia have a normal morphology in both Dmrta2 cKO and controls.

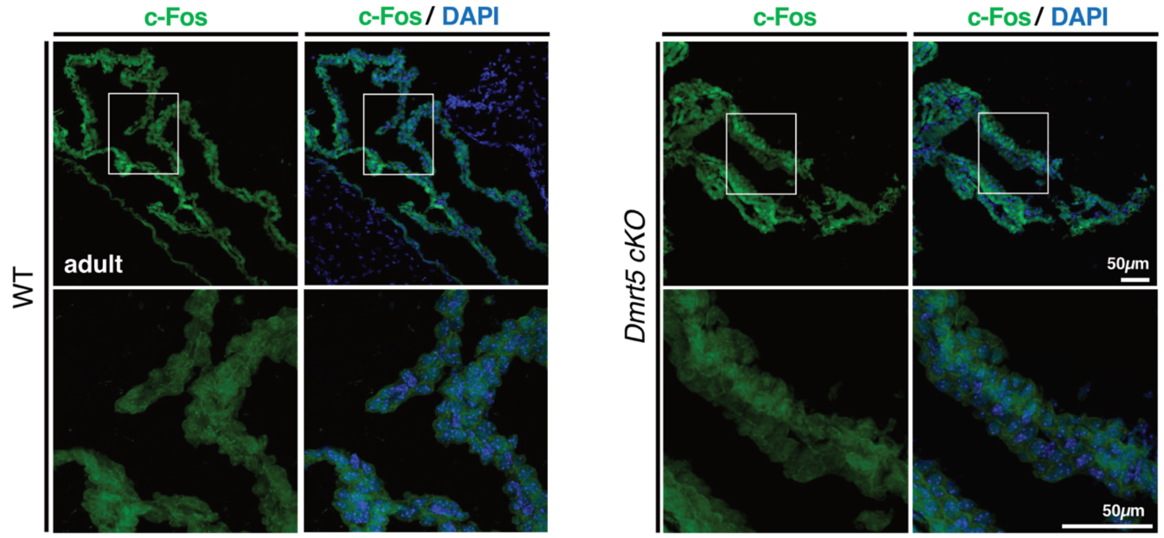


**Extended Data Figure. 7-4. $Ca^{2+}$ signalling as revealed by c-Fos immunostaining appears unaltered in the choroid plexus of *Dmrta2* cKO mice**. c-Fos immunostaining of DAPI-stained cross sections through the telencephalic choroid plexus tissue of *Dmrta2* cKO and WT control mice.

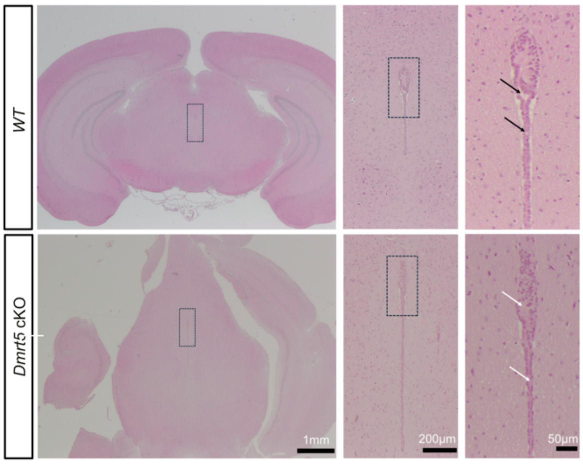


**Extended Data Figure 7-5**. **HE stained cross-sections of the brain of cKO and WT mice at the level of the thalamus showing that the aqueduct appears stenotic.** White arrows indicate levels where the passage appears obstructed. Black arrows indicate levels where the passage appears non-obstructed.
